# Fighting the spread of antibiotic resistance with bacterial competence inhibitors

**DOI:** 10.1101/683920

**Authors:** Arnau Domenech, Ana Rita Brochado, Vicky Sender, Karina Hentrich, Birgitta Henriques-Normark, Athanasios Typas, Jan-Willem Veening

## Abstract

*Streptococcus pneumoniae* is a commensal of the human nasopharynx, but it can also cause severe life-threatening antibiotic-resistant infections. Antibiotic consumption drives the spread of resistance by inducing *S. pneumoniae* competence leading to the uptake of exogenous DNA and horizontal gene transfer (HGT). We have identified potent inhibitors of competence, collectively called COM-blockers. We show that COM-blockers inhibit HGT by perturbing the proton motive force, thereby disrupting the export of the peptide that regulates competence. COM-blockers do not affect growth or compromise antibiotic activity at their active concentrations, and we did not observe any resistance development against them. Used as adjuvants of antibiotics, COM-blockers provide a strategy to reduce the spread of virulence factors and antibiotic resistance in human pathogens.

**One Sentence Summary:** Antibiotic adjuvants block competence

## Introduction

Even though *Streptococcus pneumoniae* (the pneumococcus) is part of the commensal microbiota of the human upper respiratory tract, it is also a major public health problem as it occasionally causes severe life-threatening infections, globally killing over a million people each year^1^. The alarming spread of penicillin- and multidrug-resistant *S. pneumoniae* is a cause for concern, and despite the reduction in resistant infections after the introduction of several conjugate vaccines, non-vaccine serotype clones have rapidly emerged and spread^2^. This phenomenon is especially accentuated among the at-risk population who undergo multiple courses of antimicrobial therapy per year^3^, often associated with treatment failure^4-6^.

The rapid spread of antimicrobial resistance in *S. pneumoniae* can be largely attributed to transformation, which involves the uptake and assimilation of exogenous DNA. This leads to new genotypes, and is an important mechanism of genome plasticity^7-9^. Transformation by horizontal gene transfer (HGT), the most common way DNA is acquired in *S. pneumoniae*, occurs mainly during colonization due to the simultaneous carriage of multiple pneumococcal strains^10,11^ or by the presence of closely-related *Streptococci* such as *S. mitis*, which is considered one of the major reservoirs of antimicrobial resistance and virulence genes for *S. pneumoniae*^12,13^. In contrast with other competent bacteria such as *Haemophilus influenzae or Moraxella catarrhalis*, which are able to uptake DNA during their entire life cycle, *S. pneumoniae* needs to activate a physiological state named competence to express the transformation machinery that allows the uptake and recombination of exogenous DNA (Fig. 1A)^14,15^. This is likely because pneumococci lack the canonical SOS-response and instead activate competence under stress conditions^16^. Strikingly, several commonly used antimicrobials, such as fluoroquinolones, aminoglycosides and amoxicillin-clavulanic acid, activate competence when present at sub-MIC levels^17-20^, and may thereby enhance the acquisition of virulence factors and antibiotic resistance alleles coming from the microbiota^21^. Despite the fact that antibiotics are typically administered at concentrations exceeding the MIC of the sensitive pathogen, imperfect drug penetration due to differences in the pharmacokinetics and pharmacodynamics (PK/PD) between patients, body distribution, and route of administration may contribute to local differences in final antibiotic concentration, for instance, in the nasopharynx^22^. Furthermore, local factors such as pH, the presence of species producing betalactamases, or biofilm production, can reduce antibiotic effectiveness^23^. Currently, very few studies analysed the effects of antibiotic therapy on pneumococcal carriage, but it is likely that imperfect antibiotic penetration may stimulate competence development and subsequently HGT. Indeed, a recent large-scale epidemiological study on 20,027 pneumococcal genomes showed that antibiotic resistance was associated with increased recombination^24,25^.

**Fig. 1.**
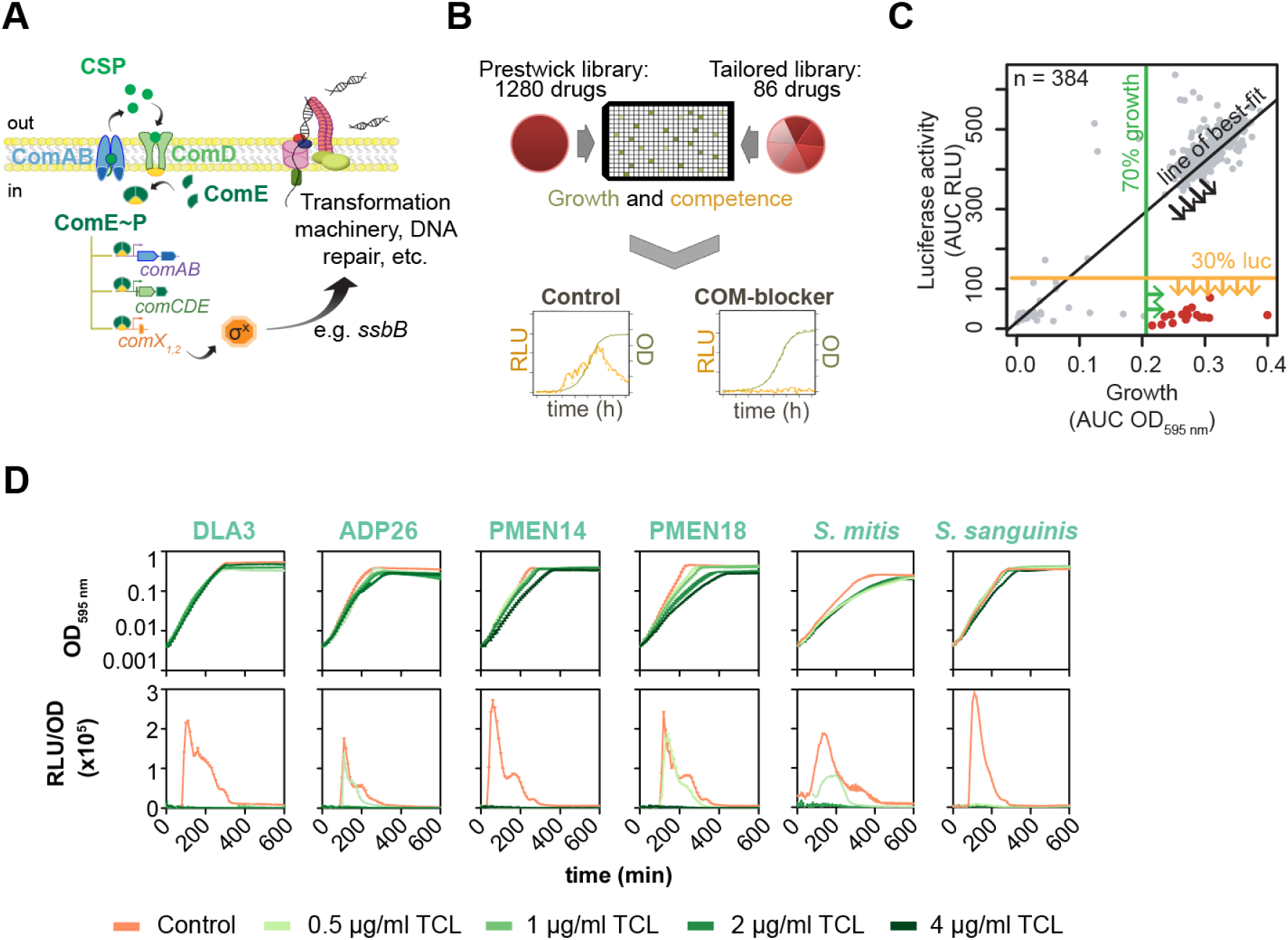
High-throughput screening to identify COM-blockers. (**A**) Overview of the regulatory network driving competence and transformation in *S. pneumoniae*. Two key operons are involved in competence activation, *comCDE* and *comAB*. The membrane transporter ComAB cleaves and exports the *comC*-encoded CSP, which then binds the membrane-bound histidine-kinase ComD, which autophosphorylates and transfers the phosphate group to the response regulator ComE. Phosphorylated ComE in turn activates the expression of the so-called ‘early’ competence genes^35^. One of them, *comX*, codes for a sigma factor (SigX), which is responsible for the activation of the so called ‘late’ competence genes, including those required for transformation and DNA repair. (**B**) Design of the COM-block screen. In control conditions (DMSO), the *S. pneumoniae* DLA3 strain (*P*_*ssbB*_*-luc*) shows a normal growth rate (green line) and activation of competence, resulting in bioluminescence (orange). Specific COM-blockers do not affect the growth rate but prevent *ssbB* promoter activity. (**C**) Example of COM-blocker identification from the high-throughput screen (data from Fig. S1A). Scatter plots show the luminescence signal (area under the curve AUC - RLU) versus growth (AUC - OD_595 nm_) obtained for each individual well per plate until 5 h of culture. The green, yellow and black lines show the stringent criteria used to identify compounds that blocked natural competence development without drastically affecting growth, which were labeled COM-blockers, here shown in red. (**D**) Growth curves and competence activity (bioluminescence) of *S. pneumoniae* variants (P_*ssbB*_*-luc*) of D39V (DLA3) and its unencapsulated variant (*cps::cat*, ADP26), two clinical strains PMEN14 (ADP49) and PMEN18 (ADP50), and two *Streptococci* of the mitis Group, *S. mitis* (ADP51) and *S. sanguinis* (ADP53), in presence of TCL. See Table S4 for more details regarding the TCL MIC of clinical strains.

In an era that antibiotic development has run dry, new strategies to preserve the current drugs and minimize the impact of resistance are required. One proposed idea is to use drug combinations^26^. For example, adjuvants with little or no antimicrobial activity on their own, can be used to promote and/or complement the antibiotic activity in different ways such as inhibiting passive resistance or targeting host defence mechanisms^27^. Such adjuvants can also go beyond the realm of common drugs to food compounds^28^. Other ideas involve searching for compounds that could reduce the evolution of resistance development, ^29,30^ or the use of quorum sensing analogous as modulators of pathogenic behaviours and players in biofilm formation^31^. Here, we propose the use of co-adjuvants to prevent transformation and the associated spread of resistant genes and virulence factors by inhibiting pneumococcal competence. We developed a high-throughput screen to identify these inhibitors, termed COM-blockers, and tested several for their ability to aid antibiotics without affecting overall treatment. We establish that the mechanism of action of COM-blockers is to disrupt the proton-motive force. This new class of small-molecule inhibitors can function as effective adjuvants to current and future antibiotic therapies, extending the lifetime of these existing treatments.

## Results

### Identification of COM-blockers

Synthetic peptide analogues that competitively inhibited specific competence-stimulating peptides (CSP)s were previously identified, and shown to reduce competence^32^. However, several CSP pherotypes exist^33^, so individual CSP inhibitors will not cover the entire CSP polymorphism spectrum. Furthermore, peptides typically do not have favourable PK/PD characteristics^34^, and it is not clear if in the nasopharynx environment where bacteria are embedded within biofilms, the CSP analogues (peptides of 17 amino acids and positively charged) are able to penetrate and interact with their target, ComD. To identify small-molecule, FDA-approved agents with favourable PK/PD characteristics that block pneumococcal competence, we screened a compound library of 1,280 drugs including antimicrobials (http://www.prestwickchemical.com), (Table S1). Given that the pneumococcal minimum inhibitory concentration (MIC) of most antibiotics in the library (∼200) is below the tested concentration (20 µM), this setup inevitably prevents us from detecting the “competence-blocking ability” of most relevant antibiotics due to strong growth defects. To overcome this drawback, we additionally tested a self-assembled library of 86 compounds (tailored library, Table S2), mostly including clinically relevant antibiotics as well as biocides that were not represented in the initial library, and selected 6 pneumococcus-tailored drug concentrations over a range of 2-fold dilutions (Table S2). In total, we tested 1,366 compounds for their ability to inhibit competence development in *S. pneumoniae* (Fig. 1B). As a reporter for competence, we used the competence-specific induced *ssbB* promoter fused to firefly luciferase (strain DLA3; *P*_*ssbB*_-*luc*)^17-20^. Luciferase activity as well as optical density were regularly measured along the growth, to determine the individual effect of each drug on competence over 5 hours. COM-blockers were identified as those compounds that strongly inhibited luciferase activity without a pronounced negative effect on growth (Fig. S1A, Materials and Methods). All experiments were done in duplicate and had high replicate correlation – average Pearson R > 0.95 (Fig. S1B). In total, we identified 46 compounds (COM-blockers; Table S3) that inhibited competence at sub-inhibitory concentrations (below MIC_90_). To exclude the possibility that COM-blockers somehow interfere with luminescence detection, for instance, reducing the amount of substrate entering the cell, we tested the effects of COM-blockers on a strain that constitutively expresses luciferase (Fig. S2A). This showed that COM-blockers do not affect luciferase production and detection, and instead specifically reduce P_*ssbB*_-*luc* expression (Fig. S2A). Two main groups of COM-blockers could be classified based on their known therapeutic activity and presumed mode of action: 20 compounds are known to affect membrane and/or ion homeostasis (Group 1), 10 are classified as antipsychotic drugs (Group 2) and the rest do not fall in these two groups (Group 3). We performed validation experiments (independent of the high-throughput screen) with 7 COM-blockers belonging to the 3 groups and confirmed strong competence inhibition (Figs. 1D and S2B). From this initial screen, we describe results for the biocide triclosan (TCL) and the antimalarial drug proguanil hydrochloride from Group 1 and pimozide from Group 2 as examples for further experiments, because they are well-studied compounds, and showed very potent COM-blocking activities at low concentration (16-32X lower than the MIC, Figs. 1D, 2A and S2).

**Fig. 2.**
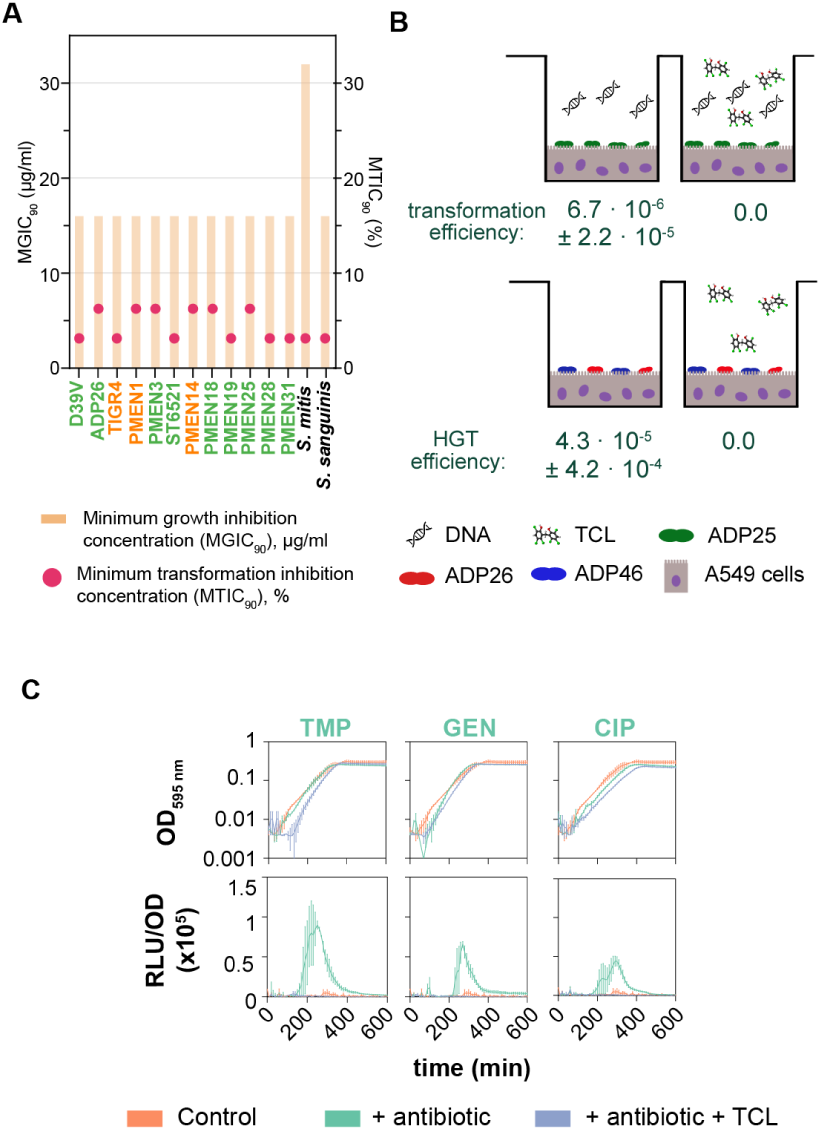
TCL effect on inhibition of DNA uptake *in vitro* in an epithelial colonisation model. (**A**) Percentage of the TCL MGIC_90_ required for inhibition of transformation (MTIC_90_) in several clinical strains and two closely related *Streptococci*. Green labelled strains express pherotype CSP-1 while orange labelled strains express pherotype CSP-2. TCL is able to inhibit natural competence in all strains, independent of the pherotype. (**B**) Upper, unencapsulated D39V bacteria (strain ADP25, green) adhered to a monolayer of A549 human lung epithelial cells (purple) were able to take up naked DNA under permissive conditions (pH 7.5). However, the addition of TCL at 2 µg/ml completely blocked transformation. Lower, HGT efficiency between two identical unencapsulated strains adhered to a monolayer of A549 human cells. ADP26 and ADP46 differ only in the presence of a resistance marker integrated in different regions of the genome (tetracycline and kanamycin, respectively). The presence of 2 µg/ml of TCL completely blocked HGT between both strains. (**C**) Unencapsulated D39V (strain ADP26) was grown in C+Y medium at pH 7.3, not permissive for natural competence development (orange). Growth curves (upper) and competence detection (lower) in presence or absence of sub-MIC concentrations of antibiotics and TCL were tested in the A549 human cells model of adhesion. Addition of antibiotics resulted in competence development (green lines); however, the presence of TCL counteracts antibiotic-induced competence (blue lines). Concentrations and antibiotics used: 0.2 µg/ml trimethoprim (TMP), 8 µg/ml gentamycin (GEN), 0.4 µg/ml ciprofloxacin (CIP), and 1 µg/ml TCL.

### TCL COM-blocker inhibits competence in clinical multi-drug resistant strains

To test whether TCL also blocks competence in other clinically relevant strains, we first examined an unencapsulated strain (ADP26), as the human nasopharynx is frequently colonized by non-typeable pneumococci characterized by a lack of capsule and higher HGT efficiencies^36^. As shown in Fig. 1D, competence was similarly inhibited in strain ADP26, although this strain was slightly less susceptible to TCL at the lowest concentration. Next, we tested several multi-drug resistant pneumococcal strains belonging to worldwide PMEN lineages^37^ with similar results (Fig. 1D). Finally, we analysed the effects of TCL in two closely-related *Streptococci* belonging to the mitis Group: *S. mitis* and *S. sanguinis*, which are abundant in the human microbiome and considered reservoirs of antibiotic resistance genes and virulence factors^12,13^. Both *S. mitis* and *S. sanguinis* demonstrated similar competence inhibition profiles compared to pneumococcal strains (Fig. 1D).

While the expression of *ssbB* is a good indicator of competence^17-20^, we wanted to examine if COM-blockers such as TCL indeed inhibit the downstream process of transformability. We first tested whether TCL would efficiently prevent transformation when naked DNA carrying relatively short homology regions (approximately 1 kb) was provided directly to the growth medium (Table S5). Secondly, we tested the transfer of a replicative plasmid (Table S5), and finally, we co-cultured two different pneumococcal strains (with a different antibiotic-resistant determinant inserted at different chromosomal positions) allowing competence-dependent HGT to take place via the uptake of chromosomal DNA (Table S6). A drastic inhibition of transformation was detected in the presence of TCL in all three experiments (Tables S5 and S6). The total number of cells recovered was similar under all the conditions, showing that the addition of TCL at competence-blocking concentrations did not affect cell viability. These results were reproduced in the presence of ciprofloxacin and kanamycin (Table S5), two antibiotics that promote competence^17,19^. Importantly, TCL also potently prevented the acquisition of exogenous DNA in clinical strains (Fig. 2B and Table S4). All clinical strains showed an MIC_90_ for growth (MGIC_90_) of 16 µg/ml TCL, with exception of *S. mitis* (32 µg/ml TCL). We here named MGIC_90_ to differentiate growth from the minimal transformation inhibition concentration (MTIC_90_), defined as the lowest concentration to inhibit the uptake of exogenous DNA in 90% of the population. Overall, 1 µg/ml of TCL (3.13 – 6.25 % of the MGIC_90_) was enough to reach MTIC_90_ in several clinical strains (Fig. 2A). Interestingly, TCL blocked HGT in clinical strains expressing two different CSP types, CSP_1_ and CSP_2_ (green and orange, respectively; Fig. 2A). This suggests that COM-blockers act by blocking a common pathway and would block DNA uptake in any pneumococcal strain, no matter the CSP allele. In total, our results establish that the presence of sub-inhibitory concentrations of TCL drastically blocks pneumococcal competence and subsequent genetic transformation, even in the presence of antibiotics that normally induce competence are present in the medium. To test whether TCL also inhibits competence development under conditions more closely resembling human colonization, we performed experiments in a human lung epithelial cell colonisation model. For these experiments, planktonic bacteria were washed to ensure that transformation and HGT occur only in bacteria colonizing the surface of the human A549 cells. To increase the natural transformation and HGT efficiency, we used unencapsulated pneumococci (ADP25, ADP26 and ADP46), since the absence of the capsule enhances adhesion thereby increasing HGT^38^. As shown in Tables S5 and S6, transformation and horizontal gene transfer was also blocked in this adherence model. Some antimicrobials induce competence at sub inhibitory concentrations^17-20^, and thereby increase the risk of developing or spreading resistance by acquiring and disseminating genetic determinants for antimicrobial resistance^8^. Importantly, we demonstrate that in combination with such competence-activating antimicrobials, TCL effectively blocks competence activation induced by antibiotic stress in a colonisation model (Fig. 2C).

A competence-inhibiting adjuvant to commonly used antimicrobials should not be cytotoxic on its own or in combination with the antimicrobial compound. To evaluate whether the co-presence of TCL plus the antibiotic could increase cytotoxicity, we examined the viability of A549 human lung epithelial cells treated with antibiotic alone and in combination with TCL. Importantly, the combination of TCL plus three common antibiotics that induce competence (ciprofloxacin, gentamycin and trimethoprim) did not show additive or synergistic toxicity when compared to the antibiotic alone after 8 h or 24 h (Fig. S3A).

As we propose to use TCL as adjuvant of clinical antibiotics, we performed checkerboard experiments (8×8 concentrations) of TCL plus several clinically relevant antibiotics to test that TCL does not compromise antibiotic activity against *S. pneumoniae*. Indeed, TCL does not interact with the activity of betalactams (cefotaxime), fluoroquinolones (ciprofloxacin), macrolides (azithromycin) and lincosamides (clindamycin) (Fig. S3B). Contrarily, a slight decreased activity of aminoglycosides was observed when 3.5 µg/ml of TCL was used (Fig. S3B). Nevertheless, this effect did not interfere with the COM-blocking activity of TCL (Fig. S3C). Furthermore, we could also exclude an interaction between TCL and clinically relevant antibiotics, including aminoglycosides, in another important human pathogen, *Pseudomonas aeruginosa*, which is intrinsically tolerant to TCL (MIC > 2 mg/ml) due to the presence of FabV, a TCL-resistant enoyl-ACP reductase^39^ (Fig. S3D).

### Undetectable resistance towards COM-block activity

We next set out to confirm whether TCL could be safely used for prolonged periods of time without losing its COM-blocking activity. Wild-type strain D39V was continuously exposed to increasing concentrations of TCL. After 30 days of consecutively plating eight independent lineages in three concentrations of TCL (10 µg/ml, 15 µg/ml and 20 µg/ml), the maximum concentration where we detected growth was 15 µg/ml of TCL, though the cells were not able to grow during continuous exposure at this concentration (Fig. 3A). This lack of resistance development might be due to TCL having multiple targets and pneumococcus lacking the main TCL target, the enoyl-acyl carrier protein (ACP) reductase FabI, present in other bacteria such as *E. coli* or *Staphylococcus aureus* (pneumococcus has a isozyme of FabI named FabK which does not interact with TCL)^40^. Importantly, 1 µg/ml of TCL could still block competence in all lineages throughout the 30 days (Fig. 3B), indicating that loss of COM-blocking activity is also undetectable even after prolonged exposures. A recent study showed that *S. aureus* and *E. coli* can develop tolerance to certain clinical antibiotics when they are previously exposed to MIC concentrations of TCL^41^. Despite the fact that our COM-blockers act at sub-inhibitory concentrations, we tested whether *S. pneumoniae* continuously exposed to TCL (lineages 1, 2, and a mix of all 8 lineages from Fig. 3A) developed reduced susceptibility to clinically relevant antibiotics. As shown in Table S7, the MICs to the tested antibiotics were constant over time showing that TCL does not promote cross-resistance towards antibiotics in *S. pneumoniae*. Finally, we also did not detect rapid resistance development against COM-blockers TCL, proguanil hydrochloride and pimozide in a standard fluctuation assay, while rifampicin resistant mutants were readily selected for in this assay (Tables S8 and S9).

**Fig. 3.**
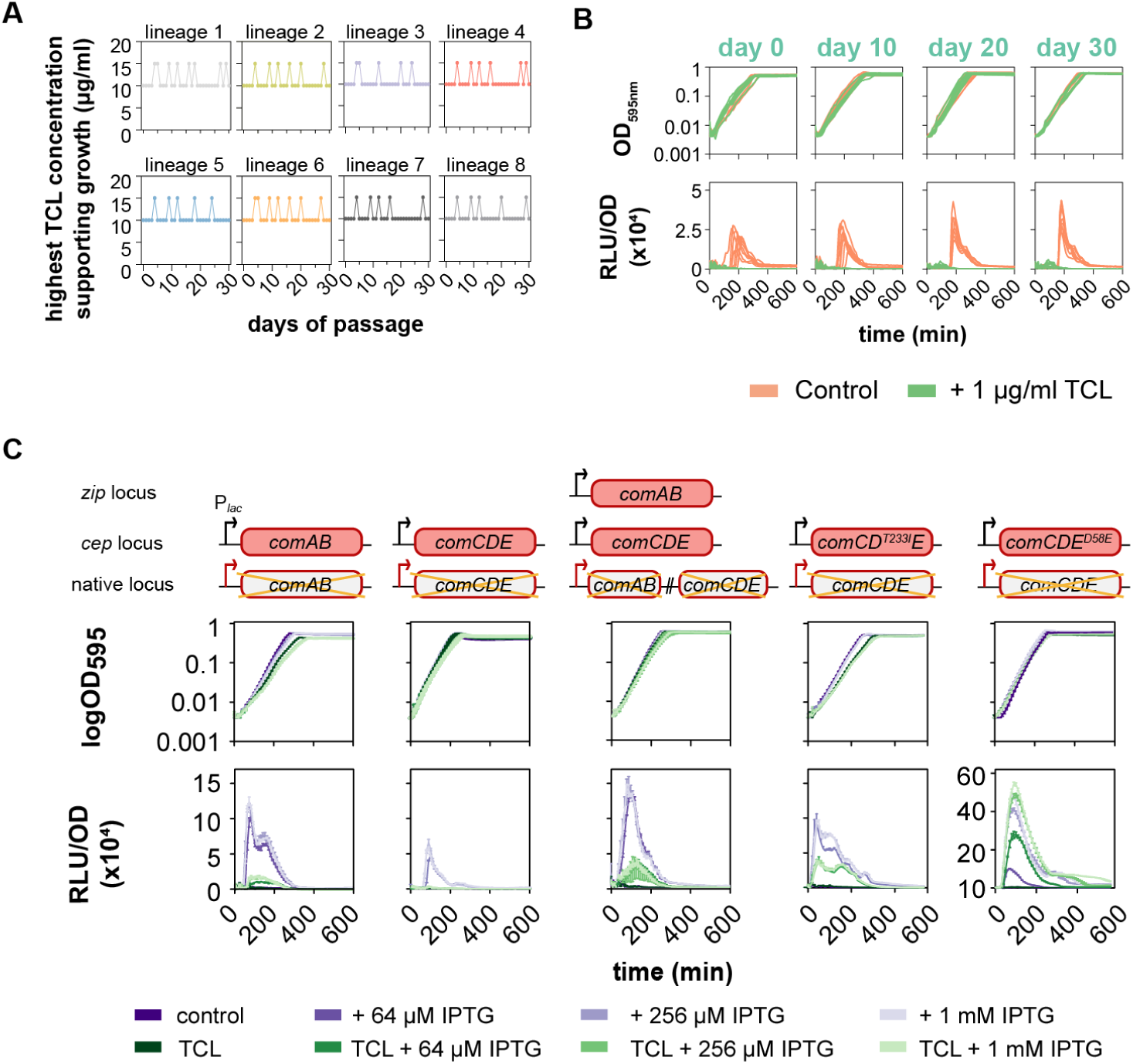
TCL blocks competence in several strains, with undetectable resistance development and no reduction in COM-blocking activity. (**A**) Undetectable acquisition of TCL resistance after 30 days of passaging in three concentrations of TCL (10 µg/ml, 15 µg/ml and 20 µg/ml TCL). Eight independent lineages were followed (one plot per lineage). For each lineage, ten different colonies were picked. Then, bacteria were plated in freshly prepared agar plates with the indicated concentrations of TCL. Only the highest TCL concentration where cells grew are indicated. The highest concentration where we detected growth was at 15 µg/ml of TCL; however, cells were not able to grow during continuous exposure at this concentration. (**B**) Continuous exposure to TCL does not affect the ability to block competence. The pool of colonies from each individual lineage was grown in C+Y medium in permissive conditions for natural competence development (pH 7.5) in the presence or absence of 1 µg/ml of TCL. (**C**) Growth curves and detection of competence development was performed in C+Y medium at pH 7.5, permissive for natural competence. IPTG was added to the medium at the beginning at different final concentrations. Strains used (left to right: ADP226, ADP107, ADP350, ADP272 and ADP148). Black arrows refer to the IPTG-inducible promoter P_*lac*_, while red arrows indicate native promoters.

To examine whether upregulation of the early operons *comAB* and/or *comCDE* could bypass the COM-blocking activity of TCL, we constructed a suite of strains in which we could ectopically induce the competence genes by adding IPTG (Fig. 3C). Upregulation of *comAB* resulted in rapid competence development; however, in the presence of TCL, competence development was still nearly abolished. Inducing expression of the entire *comCDE* operon also led to competence activation, but to a smaller extent compared to ComAB overproduction (Fig. 3C), likely because ComAB is rate-limiting in the development of competence^42^. In all cases, TCL counteracted the induction of *comAB* or *comCDE* and abolished competence (Fig. 3C). Similar effects were observed when both the *comAB* and the *comCDE* operons were induced simultaneously and with other COM-blockers such as proguanil and pimozide (Figs. 3C and S4), indicating that these competence blocking strategies are not easy to resolve and develop resistance to by raising mutations that will upregulate different competence pathways.

### Competence inhibition occurs by disrupting the proton-motive force

To identify the molecular mechanism underlying the inhibition of competence, we analysed the physiological effects of the COM-blockers on pneumococci. The slight antagonism of TCL with aminoglycosides suggested that COM-blockers could inhibit competence due to an alteration of the proton-motive force (PMF), as aminoglycosides require active electron transport^43^. Indeed, classical disrupters of the PMF (carbonylcyanide m-chlorophenylhydrazone [CCCP], nigericin and valinomycin) were also highly effective in blocking competence (Fig. S2B).

Next, we tested the flux of H^+^ and K^+^ after addition of several COM-blockers to understand how they perturb the PMF (Fig. 4A-B). TCL decreases the internal pH, but, in contrast to nigericin, TCL and proguanil also decrease intracellular potassium levels. Interestingly, pimozide did not affect the internal pH but did affect K^+^ uptake. This suggests that targeting the PMF is promising for blocking competence development, independent of the underlying mechanism of its disruption. To confirm that TCL decreases the intracellular pH, we tested the susceptibility of *S. pneumoniae* to TCL in a range of pH values. Indeed, the stepwise acidification of the medium by decreasing the pH resulted in increased susceptibility to TCL (Fig. 4C).

**Fig. 4.**
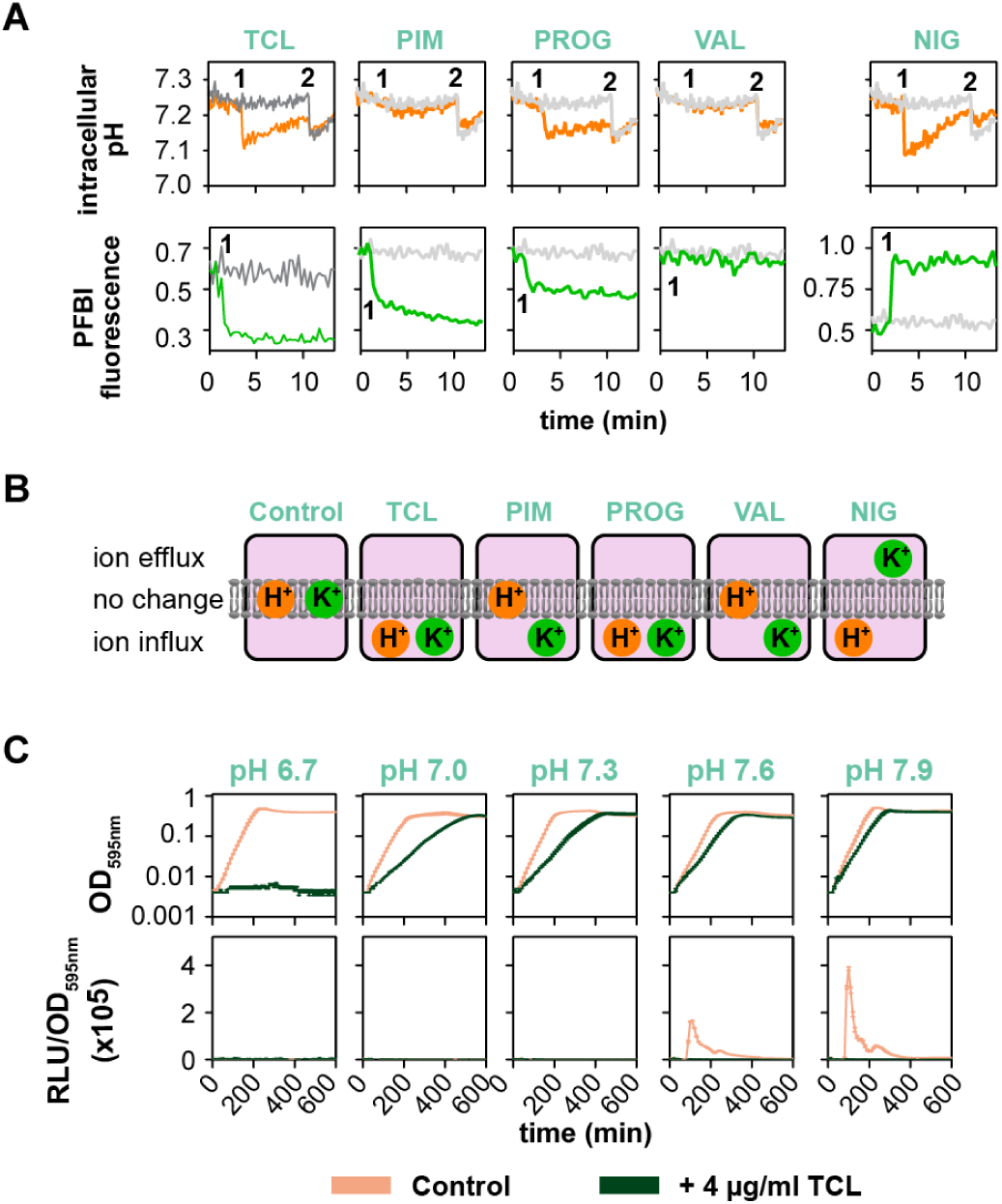
COM-blockers disrupt the proton-motive force (PMF). (**A**) Intracellular changes in pH (H^+^) and potassium (K^+^) after the addition of several COM-blockers: 4 µg/ml of TCL, pimozide (PIM) and proguanil hydrochloride (PROG), or 0.5 µg/ml of NIG and VAL. NIG, an antiporter of H^+^ and K^+^, decreases the internal pH by exchanging H^+^ and potassium^43^. VAL makes selective pores for K^+^. Upper panels; pneumococci were loaded with the pH-sensitive dye BCECF (50 µM). After the baseline was recorded, TCL (orange) or DMSO (grey) were added (1). In presence of TCL, PROG and NIG, a drop in signal was observed, indicating a decrease in the internal pH. The addition of 20 µg/ml of NIG (2) collapses the PMF and equilibrates the external and internal pH. Lower panels; cells were incubated with PBFI/AM dye (5 µM) and treated (1) with COM-blockers (green) or DMSO (grey). Changes in fluorescence were recorded, showing that TCL, PIM, PROG lead to increased intracellular accumulation of K^+^. Contrary, NIG addition results in an extrusion of K^+^ outside the cell. (**B**) Summary of the effect on internal pH and potassium levels by the addition of several COM-blockers. TCL decreases the internal pH but also increases the cytoplasmic concentration of K^+^. (**C**) Effect of TCL on growth (upper) and competence activity (lower) of *S. pneumoniae* DLA3 grown in a range of pH values. Acidic medium increases the susceptibility of DLA3 to 4 µg/ml of TCL, with complete growth abolition at pH 6.7 and lower

To confirm whether pH homeostasis is essential for competence, we targeted the F_0_F_1_ proton ATPase complex, which in *S. pneumoniae* is used to maintain pH homeostasis^44^. We first used proguanil hydrochloride, an antimalarial drug that is known to directly inhibit the pneumococcal F_0_F_1_ ATPase^45^ and was also found to block competence in our initial high-throughput screen (Figs. 1B and S2B)^45^. We additionally used CRISPR interference (CRISPRi)^45^ to reduce transcription of *atpE, atpA* and *atpC*, three genes of the operon encoding the ATP synthase (Fig. S5A). Indeed, the reduced expression of the *atp* genes (Fig. S5B) led to a strong inhibition of competence.

To confirm that the absence of bioluminescence was due to the block in competence and not to the absence of luciferase activity (as this enzyme requires ATP to convert luciferine to oxyluciferine), we repeated the same experiments using a constitutive reporter. In this case, we did not observe significant differences (Fig. S5C).

To strengthen the case for perturbations in the PMF causing COM-blocking effect, we searched for evidence of other COM-blockers that also inhibit the F_0_F_1_ proton ATPase enzyme. One well known compound, optochin (ethylhydrocupreine hydrochloride), is an antibiotic classically used to differentiate *S. pneumoniae* (optochin-susceptible) from other α-haemolytic streptococci (optochin-resistant)^47^. However, pneumococcal optochin resistance has been reported by point mutations in *atpE*, a subunit of the F_0_F_1_ proton ATPase enzyme^48^. We first confirmed that optochin is a COM-blocker (Fig. S5D) and then tested whether these two different amino acid changes in AtpE (AtpE^V48L^ and AtpE^A49T^) led to COM-blocker resistance. As expected, both mutations conferred resistance to optochin, which resulted in a loss of COM-blocking activity (Fig. S5D). However, there was no loss of COM-blocking by TCL or proguanil in AtpE mutants, since neither TCL nor proguanil are expected to have direct interaction with the ATPase (Fig. S5E-F). In total, we conclude that COM-blockers work by perturbing the PMF.

### COM-blockers perturb CSP export

For a better understanding of how PMF disruption inhibits competence, we tested if COM-blockers alter any of the known membrane processes essential for competence development: the secretion of ComC (called CSP once outside the cell) by the ComAB exporter and/or the ability of ComD to phosphorylate ComE (Fig. 1A). First, we evaluated if the presence of TCL affected the interaction between ComD and ComE by adding exogenous CSP_1_ at different times to the control strain (DLA3) and a *comAB* mutant (ADP342). Shortly after the addition of CSP_1_, a rapid bioluminescence signal was detected in both strains (Fig. 5A); however, competence was switched off earlier in the *comAB* mutant, probably because this strain cannot amplify the competence signal by exporting extra CSP_1_ in the common pool (Fig. 5A).

**Fig. 5.**
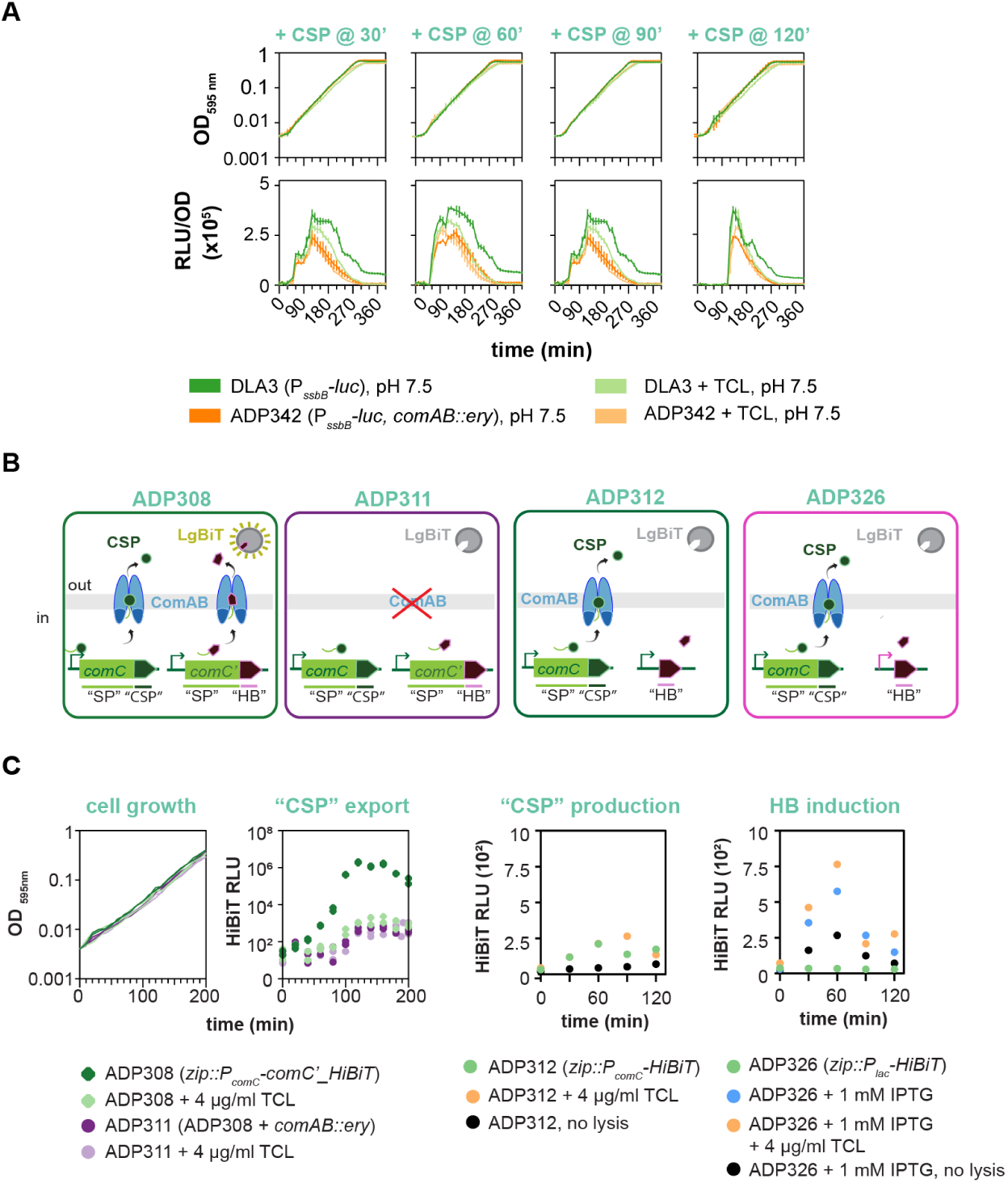
Triclosan (TCL) affects CSP export. **(A)** Competence blocking by TCL is bypassed by addition of synthetic CSP_1_. 100 ng/ml CSP_1_ was added to the medium at the indicated times: after 30’, 60’, 90’ and 120’ to strains DLA3 (*P*_*ssbB*_*-luc*) and ADP342 (*P*_*ssbB*_*-luc, comAB::ery*). TCL was present at 4 µg/ml instead of 1 µg/ml to exacerbate the phenotype. (**B**) Graphical representation of the HiBiT experiments. From left to right: (I) ComC (called CSP once outside the cell) and HiBiT are regulated by the *comCDE* promoter, and both precursors have a signal peptide sequence that is recognized, cleaved and exported by ComAB (strain ADP308). Once outside the cell, HiBiT interacts with the soluble protein LgBiT and produces bioluminescence. (II) In the absence of ComAB (strain ADP311), HiBiT accumulates in the cytoplasm since it cannot be recognized and exported. (III and IV) HiBiT peptide expression is under control of P_*comCDE*_ promoter (ADP312) or under control of IPTG-inducible P_*lac*_ promoter (ADP326) without the leader sequence. In both cases, HiBiT is accumulated in the cytoplasm since it cannot be recognized and exported. (**C**) CSP export is strongly reduced in presence of TCL. Bioluminescence (relative luminescence units, RLU) can be correlated with CSP export. Left, CSP is exported until the saturation point (∼10^7^ RLU). The presence of TCL abolished CSP export, showing similar background RLU to the *comAB* mutant strain (ADP311). To confirm that TCL does not affect the expression of the HiBiT, ADP312 and ADP326 were grown in C+Y at pH 7.9 and samples were taken every 30 min for analysis and were lysed with 0.1% triton X100 and 2% deoxycholate. pH 7.9 used in this experiment allowed earlier competence development compared to the pH used in previous experiments (pH 7.5), thereby reducing the required amount of substrate and number of reads to favour a rapid natural competence.

Furthermore, the addition of CSP_1_ to cells treated with TCL also showed rapid competence activation similar to the control, indicating that ComD can recognize CSP and phosphorylate ComE in the presence of TCL. However, as observed for the *comAB* mutant, the wild-type DLA3 strain in the presence of TCL showed faster competence shut down, suggesting that TCL inhibits either CSP production or export (Fig. 5A). In line with the observations that TCL decouples the CSP-ComD-ComE positive feedback (Fig. 1A), the addition of TCL strongly reduced the activity of both P_*comAB*_ and P_*comCDE*_. This potentially decreases the amount of CSP accumulated and thereby leads to a loss of competence development (Fig. S6).

To confirm if CSP production and/or export are inhibited by TCL, we employed a split-luciferase HiBiT-tag detection system to track the CSP export rate^20^. When HiBiT is expressed and exported out of the cell, it reacts with a HiBiT-dependent luciferase variant (LgBiT) added to the medium, resulting in bioluminescence (Fig. 5B). First, the HiBiT tag was placed under the control of the *comCDE* promoter and coupled to the signal peptide sequence of *comC* (ADP308). When HiBiT is produced in this system, ComAB recognizes the leader signal, cleaves it and exports HiBiT. In the absence of ComAB (in strain ADP311), HiBiT is not matured and exported, so it accumulates in the cytoplasm and no luminescence is generated (Fig. 5C, left). In the presence of TCL, bioluminescence was abolished, indicating that HiBiT, and thereby CSP, were either not produced or were not exported (Fig. 5C). Finally, in the absence of the signal sequence (strain ADP312), HiBiT accumulated in the cytoplasm, and luminescence was not generated unless cells were lysed (Fig. 5C).

The results presented so far suggest that COM-blockers such as TCL prevent the export of CSP, thereby blocking competence. Indeed, it is reasonable to suggest that the PMF influences the activity of peptidase-containing ATP-binding cassette transporters such as ComAB, as they require ATP to transport substrates^49^. If this is the case, then the ectopic expression of a constitutively active mutant of ComD or ComE should be insensitive to TCL. As the overproduction of ComE or ComD alone repressed competence because unphosphorylated ComE represses ComE-dependent promoters (Fig. S7)^50^, we constructed two strains in which the entire *comCDE* operon was under IPTG-inducible control, but in which either the wild-type *comD* or *comE* genes were exchanged for two constitutively active mutants. The first mutant, ComD^*T233I*^ (strain ADP272), constitutively phosphorylates ComE independent of the presence of CSP^30^, while the second, ComE^D58E^ (strain ADP148), mimics the phosphorylated form of ComE, thus inducing competence even in the absence of CSP or ComD^35^. ComD^*T233I*^ showed a hypercompetent phenotype in presence of IPTG, though TCL was still able to reduce competence activation to nearly half the luminescence production when ComD^*T233I*^ was induced (Fig. 3C). The differences between the presence or not of TCL, suggests that the competence feedback is still necessary to fully activate this strain, and the export of CSP boosts the positive feedback increasing the bioluminescence signal, or that a fully intact PMF is required for ComD-ComE signalling. In contrast, competence was unaffected by TCL when ComE^D58E^ was induced.

Taken together, we conclude that PMF disruption by COM-blockers affects CSP export. Consequently, ComD will not autophosphorylate and will not phosphorylate ComE, thereby also reducing P_*comAB*_ and P_*comCDE*_ transcription (Fig. 6). As the competence pathway is tightly regulated at different levels to prevent spontaneous activation when it is not required, a minimal disruption on the CSP export is enough to unbalance the accumulation of ComE and preventing competence development (Figs. 6 and S6). For this reason, the here mentioned COM-blockers disrupt competence without attenuating cell growth, as they already work at sub-inhibitory concentrations (16-32% below the MIC).

**Fig. 6.**
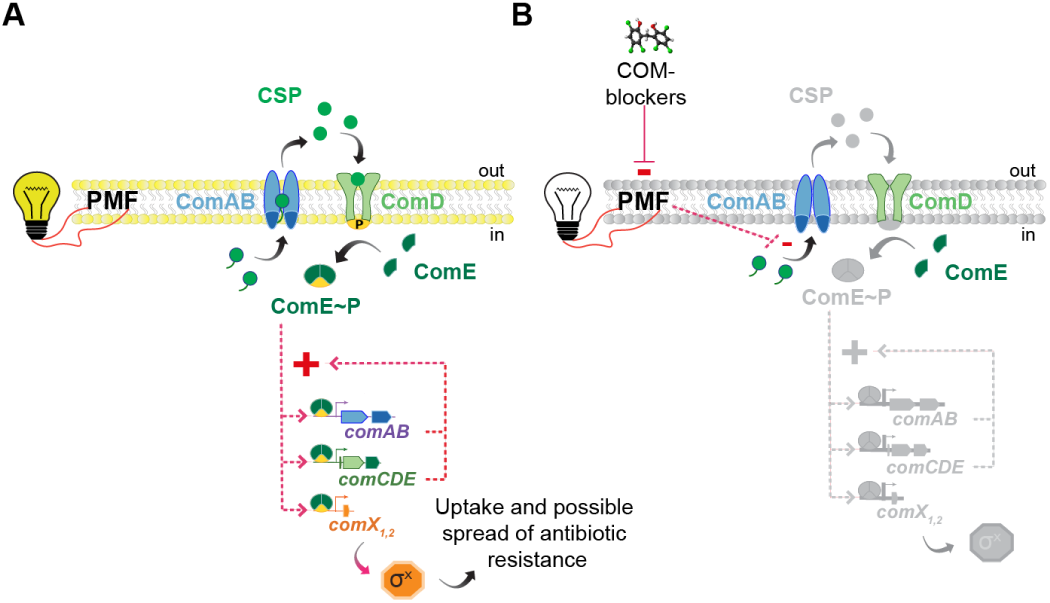
Model of COM-blockers activity. **(A)** The Proton Motive Force (PMF) maintains membrane homeostasis and thus, membrane-associated processes such as ComAB activity. ComAB cleaves and exports the *comC*-encoded competence-stimulating peptide (CSP). CSP then stimulates the phosphorylation of ComE via ComD (see Fig. 1A for more details), creating a positive feedback loop, amplifying the signal (by increasing *comAB* and *comCDE* expression) and inducing the expression of *comX*, which encodes for the sigma factor SigX, responsible for the activation of the transformation machinery, which can lead to the acquisition of antibiotic resistance. **(B)** COM-blockers disrupt the PMF, thereby preventing ComAB from exporting enough CSP to trigger the positive feedback (grey pathway). The absence of feedback results in a decreased expression of both *comAB* and *comCDE* operons, and the absence of competence and the expression of the transformation machinery.

## Conclusion

The COM-blockers developed herein represent a new class of treatment in the fight against antibiotic resistance in which they themselves do not eliminate infections, but instead assist antibiotics to prevent the acquisition of resistance genes. Importantly, COM-blockers are already FDA approved and some of them, such as proguanil hydrochloride, has been used safely as an antimalarial medication since its approval in 2000 (https://www.accessdata.fda.gov). Future studies should be aimed at assessing optimal dosing in combination with antibiotics and safety studies. The application of COM-blockers as small-molecule adjuvants to enhance the activity of existing antibiotics could translate into new therapies that will extend the time both new and old antibiotics can be used before the arrival of wide-spread resistance. Moreover, they could also act as “anti-evolution” drugs that, in combination with the antibiotic treatment, could prolong the use of the current conjugate vaccines and reduce serotype displacement.

## Materials and Methods

### Bacterial strains, human cells and growth conditions

Bacterial strains used in this study are listed in Table S10. Growth conditions of bacterial cells were described previously17. Briefly, S. pneumoniae was grown in C+Y medium (pH 6.8, non-permissive conditions for natural competence induction), at 37°C and stored at −80°C in C+Y with 14.5% glycerol at OD595 nm of 0.4. Determination of minimum inhibitory concentration (MIC) and minimum bactericidal concentration (MBC), was performed following the Clinical Laboratory Standards Institute (CLSI) methods51.

The cell line A549 (human lung carcinoma cell line) was used for in vitro experiments as follows: cells were seeded in 96-well plates at a density of 1.5·106 cells/cm2 and maintained in GlutaMAX(tm) media (Gibco®) at 37°C + 5% CO2.

### Competence assays

The *S. pneumoniae* strains were cultured in a Tecan Infinite F200 PRO allowing for real-time monitoring of competence induction *in vitro. S. pneumoniae* was grown to OD_595_ 0.4 in C+Y pH 6.8 +/- 0.05 (non-permissive conditions for natural competence induction) at 37°C. This pre-culture was diluted 100-fold in C+Y pH 7.5 +/- 0.05 (permissive conditions for natural competence induction) containing 0.45 mg/ml of luciferine and then incubated in 384 or 96-wells microtiter plates with no shaking. Growth (OD_595 nm_) and luciferase activity (RLU) were measured every 10 min during 14h, with at least three replicates per condition, except for high-throuput screening where we did duplicates. Expression of the *luc* gene (only if competence is activated) results in the production of luciferase and thereby in the emission of light^17^. An average of 3 replicates and the standard error of the mean (SEM) are shown unless indicated.

### High Throughput Screen (HTS) of inhibitors of competence in *S. pneumoniae*

The Prestwick library was tested against the DLA3 (P_*ssbB*_*-luc*) strain, in order to identify potential competence inhibitors. This library consists of 1280 FDA- and EMA-approved compounds (arranged in 4×384-wells microtiter plates), including a broad variety of antimicrobials and other drugs covering a wide range of chemical structures and therapeutic effects (http://www.prestwickchemical.com). Competence and growth assays were done simultaneously, in duplicate, in 384-wells microtiter plates containing 100 µl as described above, with a unique concentration of 20µM for all compounds. The self-assembled pneumococcus tailored library containing 86 relevant clinical antimicrobials and several biocides in a range of 6 serial dilutions was tested using an identical setup. The library was assembled in 1×384-wells microtiter plate, with 4 concentrations per drug, over a 2-fold serial dilution. Starting concentrations of all individual compounds are listed on Table S1. This library was tested twice – once at the concentrations described in Table S1, and a second time with ¼ of those. This enabled us to expand the concentration range per drug from 4 to 6 events, along 2-fold serial dilutions. We obtained duplicates for both cases, full & ¼ initial drug concentration. All liquid handling involving drugs and bacterial cultures was done with a Biomek FX liquid handler (Beckman Coulter). Plates were sealed with breathable membranes (Breathe-Easy®).

Here we define competence-blocking compounds – COM-blockers – as all compounds that inhibit competence as measured by luciferase activity under control of the *ssbB* promoter, and do not strongly halt growth. We used Area Under the Curve (AUC) until 5h culture-time to describe growth (OD_595 nm_) and luciferase activity/competence (RLU) per well. Data reproducibility was assessed by Pearson correlation between the 2 replicates of each library plate (4 plates Prestwick library & 2 plates tailored library). We obtained very high reproducibility across replicates: 0.95 for growth and 0.94 for luciferase activity on average across all plates (Fig. S2). The vast majority of tested compounds do not show any specific effect on luciferase activity when compared to growth – when growth is decreased, luciferase activity will also decrease accordingly (Fig. S1). COM-blockers will deviate from this rule, as they will inflict low luciferase activity, while minimally affecting bacterial growth. High confidence COM-blockers satisfy the following stringent criteria (Fig. S1): a) below 7.5% quantile of the residuals of the line of best-fit luciferase activity versus growth; b) luciferase activity is below 70% of the median luciferase activity across all compounds in the plate and c) growth is at least 70% of median growth across all compounds in the plate. Relaxing these cutoffs causes lower agreement between the 2 replicates. Data analysis was done per individual plate, in order to account for eventual batch-effects that could occur across the different plates. R version 2.15.1 & RStudio Version 1.0.136 were used.

### Checkerboard assays

In order to find possible drug-drug interactions between triclosan and the most common antibiotic classes, several dose-resolved checkerboards were performed in 384-wells microtiter plates. Combination of 8×8 linearly spaced concentrations of each drug were tested for at least two representative antibiotics of each class: B-lactams, fluoroquinolones, macrolides and aminoglycosides (antibiotic concentrations are provided with the corresponding figures). Liquid handling was done using a Biomek FX liquid handler (Beckman Coulter).

### Horizontal gene transfer (HGT) between S. pneumoniae strains

We used the DLA3 (tetracycline resistant) and MK134 (kanamycin resistant) strains, which are naturally competent and are genetically identical, with exception of the locus where *P*_*ssbB*_-*luc* was inserted, and the antimicrobial marker used (Table S5). Strains were grown to OD_595 nm_ 0.4 in C+Y pH 6.8 at 37°C (non-permissive conditions for natural competence activation). Then, a mixed 100-fold dilution of both strains were grown in C+Y pH 7.5 (permissive conditions) to OD_595 nm_ to promote the transfer of genes. Afterwards, serial dilution of cultures were plated without antibiotics (for the recovery of the total viable counts) and with the combination of 250 μg/ml of kanamycin plus 1 μg/ml tetracycline. The experiments were performed adding either 1 μg/ml or 2 μg/ml of TCL, and comparing with the control condition without TCL. Each experimental condition was independently performed at least three times.

For the replicative plasmid experiments, we used the strain PGs6 (erythromycin-resistant) as a donor, and the strain ADP65 (kanamycin- and gentamycin-resistant) as a recipient. The experiments were performed as indicated above, but plating in Columbia blood agar containing all three antibiotics (0.25μg/ml of erythromycin, 250 μg/ml of kanamycin and 20μg/ml of gentamycin). Plates were supplemented with the three antibiotics in order to nearly exclude the possibility of horizontal gene transfer from the recipient to the donor strain.

### Transformation assays

The wild-type D39V *S. pneumoniae* strain was grown to OD_595_ 0.4 in C+Y pH 6.8 at 37°C. This pre-culture was diluted 100-fold in C+Y pH 7.5 (permissive conditions) and 1 µg/ml or 2 µg/ml of TCL was added to the culture. After 150 min of incubation at 37°C (time required for natural competence induction, data not shown), 1 µg/ml of plasmid pLA18 (carrying the tetracycline-resistance determinant *tetM*^17^, was added (Table S9 for plasmid information). The cultures were incubated at 37°C for 1h30min and then serial dilutions were plated either with or without 1 µg/ml of tetracycline. Transformation efficiency was calculated by dividing the number of transformants by the total number of viable count. Three independent replicates of each condition was performed.

### In vitro MTT cytotoxicity assays

Triclosan is a drug approved for the Food and Drug Administration (FDA), which is not toxic at low concentration. Nevertheless, cytotoxic assays with a TCL concentration range in combination of several antimicrobials were performed, using the MTT assay procedure^54^. A549 cells were incubated in 96-well plates in a density of 1.5e6 cells/cm2, and grown at 37 °C with 5% CO2. A TCL range of concentrations and/or in combination with antimicrobials were added to the wells with fresh colourless culture medium (100 µl). Then, plates were incubated for 8h and 24h. At the time indicated, 0.1 ml of MTT solution (3-(4,5-dimethylthiazol-2-yl)-2,5-diphenyl tetrazolium bromide) was added to the wells following the recommendation of the commercial kit (Vybrant® MTT Cell proliferation assay kit, ThermoFisher Scientific). After 2h of incubation at 37°C, 50 µl of DMSO buffer was added for 10 min, and the optical density was measured as absorbance at 570 nm. Each experiment was repeated at least three times to obtain the mean values.

### Transformation and HGT assays in bacteria adhered to human airway cells

The same experiments previously described were reproduced in *S. pneumoniae* adhered to a monolayer of human airway cells. A549 cells were incubated in 96-well plates following the same criteria than in the MTT cytotoxicity assay. After incubation, cells were washed twice with PBS and were fixed in 4 % paraformaldehyde for 10 min. Then they were washed three times with PBS and were inoculated with a *S. pneumoniae* preculture in C+Y at OD 0.04. After 10 minutes, the 96 well plate was spun down 1 minute at 2000 r.p.m. and the supernatant was removed. Wells were washed with PBS in order to remove planktonic cells, and fresh media supplemented with Triclosan and/or antibiotics were added in order to reproduce the experiments described above.

### Nano-Glo HiBiT Extracellular Detection System

Cells were pre-cultured in C+Y (pH 6.8) at 37°C to an OD_595 nm_ of 0.1, washed and diluted as explained before in C+Y (pH 7.9). Experiments were started with an inoculation density of OD_595nm_ 0.001. Every 20 minutes, 5 µl of the Nano-Glo® Extracellular Detection System reagent was added as specified in the manufacturer’s instructions, and bioluminescence was detected in 96-wells plates with a Tecan Infinite 200 PRO luminometer at 37°C. Additionally, media and PBS samples were used as controls. Bioluminescence was measured right after the reagent addition. Two replicates for each time point and condition were tested.

### Monitoring of ions fluxes

Cultured *S. pneumoniae* ADP25 cells at OD_595 nm_ 0.4 were washed and resuspended in buffer (5 mM HEPES buffer, 20 mM glucose, 100 mM KCl, pH 7.4). The measurement of the intracellular fluxes of K^+^ and H^+^ were analysed using the appropriate dyes: PFBI, AM; and BCECF, AM; respectively, and following the protocol^55^. Valinomycin, Nigericin and CCCP were used as controls of proton motive force disturbers.

### Measuring Mutation Rates Using the Fluctuation Assay

*S. pneumoniae* D39V was grown to OD_595_ 0.4 in C+Y pH 6.8 +/- 0.05 at 37°C. Then, on the basis that OD_595 nm_ 0.1 contains 1.5×10^8^ colony forming units per ml (CFU/ml)_56_, 6×10^8^ cfu’s were plated onto Colombia blood agar, containing MIC or 2X MIC concentrations of the COM-blockers TCL, proguanil and pimozide (16 μg/ml and 32 μg/ml for each compound). Extra plates containing rifampicin (0.03 μg/ml and 0.06 μg/ml) were used as positive control for the rapid selection of mutants. Three independent biological replicates were performed.

Number of colonies were counted after 24h and 48h, and re-streaked with the same concentration of antibiotic (Table S8). Then *rpoB* of ten spontaneously resistant colonies from rifampicin treatment was analysed by Sanger sequencing with the primer CGGTGACTCCTGCAGATATCCTTGCTGAG to exclude false positive colonies (Table S9).

## Author Contributions and Notes

Conceptualization: A.D. and J.W.V; methodology: A.D., V.S., A.R.B and K.H.; writing - original draft: A.D. and J.W.V.; writing – review & editing: A.D., V.S., A.R.B, K.H., A.T., B.H.N., and J.W.V.

## Competing interests

A.D. and J.W.V. authorship the International Patent Application No. PCT/NL2017/050671 WO/2018/070874.

This article contains supporting information online.

## Acknowledgments

We are grateful to Mark van der Linden and Peter Hermans for providing the collection of clinical PMEN strains tested in this study. We thank Marco Oggioni for insightful discussions regarding the pneumococcal F_0_F_1_ ATPase and are grateful to all members of the Veening lab for stimulating discussions.

Work in the Veening lab is supported by the Swiss National Science Foundation (project grant 31003A_172861), a JPIAMR grant (50-52900-98-202) from the Netherlands Organisation for Health Research and Development (ZonMW) and ERC consolidator grant 771534-PneumoCaTChER. A.D. was supported by Marie Skłodowska-Curie fellowship 657546.

## References

1. Wahl, B. et al. Burden of Streptococcus pneumoniae and Haemophilus influenzae type b disease in children in the era of conjugate vaccines: global, regional, and national estimates for 2000-15. Lancet Glob. Health. 6, e744–e757 (2018).

2. Kim, L., McGee, L., Tomczyk, S., Beall, B. Biological and Epidemiological Features of Antibiotic-Resistant Streptococcus pneumoniae in Pre- and Post-Conjugate Vaccine Eras: a United States Perspective. Clin. Microbiol. Rev. 29, 525–552 (2016).

3. Domenech, A. et al., Dynamics of the pneumococcal population causing acute exacerbations in COPD patients in a Barcelona hospital (2009-12): comparison with 2001-04 and 2005-08 periods. J. Antimicrob. Chemother. 69, 932–939 (2014).

4. Pletz, M. W. R., McGee, L., Burkhardt, O., Lode, H., Klugman, K. P. Ciprofloxacin treatment failure in a patient with resistant Streptococcus pneumoniae infection following prior ciprofloxacin therapy. Eur. J. Clin. Microbiol. Infect. Dis. Off. Publ. Eur. Soc. Clin. Microbiol. 24, 58–60 (2005).

5. Rzeszutek, M. et al. A review of clinical failures associated with macrolide-resistant Streptococcus pneumoniae. Int. J. Antimicrob. Agents. 24, 95–104 (2004).

6. Zielnik-Jurkiewicz, B., Bielicka, A. Antibiotic resistance of Streptococcus pneumoniae in children with acute otitis media treatment failure. Int. J. Pediatr. Otorhinolaryngol. 79, 2129–2133 (2015).

7. Chewapreecha, C. et al. Dense genomic sampling identifies highways of pneumococcal recombination. Nat. Genet. 46, 305–309 (2014).

8. Croucher, N. J. et al. Rapid pneumococcal evolution in response to clinical interventions. Science. 331, 430–434 (2011).

9. Gladstone, R. A. et al. International genomic definition of pneumococcal lineages, to contextualise disease, antibiotic resistance and vaccine impact. EBioMedicine. 43, 338–346 (2019).

10. Donkor, E. S. et al. High levels of recombination among Streptococcus pneumoniae isolates from the Gambia. mBio. 2, e00040–00011 (2011).

11. Marks, L. R., Reddinger, R. M., Hakansson, A. P. High Levels of Genetic Recombination during Nasopharyngeal Carriage and Biofilm Formation in Streptococcus pneumoniae. mBio. 3, e00200–12 (2012).

12. Bryskier A. Viridans group streptococci: a reservoir of resistant bacteria in oral cavities. Clin Microbiol Infect. 8, 65–9 (2002).

13. Janoir, C., Podglajen, I., Kitzis, M. D., Poyart, C., Gutmann, L. In vitro exchange of fluoroquinolone resistance determinants between Streptococcus pneumoniae and viridans streptococci and genomic organization of the parE-parC region in S. mitis. J. Infect. Dis. 180, 555–558 (1999).

14. Claverys, J.-P., Martin, B., Polard, P. The genetic transformation machinery: composition, localization, and mechanism. FEMS Microbiol. Rev. 33, 643–656 (2009).

15. Johnston, C., Martin, B., Fichant, G., Polard, P., Claverys, J-P. Bacterial transformation: distribution, shared mechanisms and divergent control. Nat. Rev. Microbiol. 12, 181–196 (2014).

16. Charpentier, X., Polard, P., Claverys, J. P. Induction of competence for genetic transformation by antibiotics: convergent evolution of stress responses in distant bacterial species lacking SOS? Curr Opin Microbiol. 15, 570–6 (2012).

17. Slager, J., Kjos, M., Attaiech, L., Veening, J.-W. Antibiotic-induced replication stress triggers bacterial competence by increasing gene dosage near the origin. Cell. 157, 395–406 (2014).

18. Stevens, K. E., Chang, D., Zwack, E. E., Sebert, M. E. Competence in Streptococcus pneumoniae is regulated by the rate of ribosomal decoding errors. mBio. 2, e00071–00011 (2011).

19. Prudhomme, M., Attaiech, L., Sanchez, G., Martin, B., Claverys, J.-P. Antibiotic stress induces genetic transformability in the human pathogen Streptococcus pneumoniae. Science. 313, 89–92 (2006).

20. Domenech, A., Slager, J., Veening, J.-W. Antibiotic-Induced Cell Chaining Triggers Pneumococcal Competence by Reshaping Quorum Sensing to Autocrine-Like Signaling. Cell Rep. 25, 2390-2400.e3 (2018).

21. Veening, J.-W. & Blokesch, M. Interbacterial predation as a strategy for DNA acquisition in naturally competent bacteria. Nat. Rev. Microbiol. 15, 629 (2017).

22. Moreno-Gámez, S. et al. Imperfect drug penetration leads to spatial monotherapy and rapid evolution of multidrug resistance. Proc Natl Acad Sci U S A.112, E2874–83 (2015).

23. Levison, M. E., Levison, J. H. Pharmacokinetics and pharmacodynamics of antibacterial agents. Infect Dis Clin North Am. 23: 791–815 (2009).

24. Gladstone, R. A. et al. International genomic definition of pneumococcal lineages, to contextualise disease, antibiotic resistance and vaccine impact. EBioMedicine. 43, 338–346 (2019).

25. Lo, S. W. et al. Pneumococcal lineages associated with serotype replacement and antibiotic resistance in childhood invasive pneumococcal disease in the post-PCV13 era: an international whole-genome sequencing study. Lancet Infect Dis. 7, pii: S1473-3099(19)30297-X (2019),

26. Tyers, M. & Wright, G. D. Drug combinations: a strategy to extend the life of antibiotics in the 21st century. Nat Rev Microbiol. 17: 141–155 (2019).

27. G. D. Wright, Antibiotic Adjuvants: Rescuing Antibiotics from Resistance. Trends Microbiol. 24, 862–871 (2016).

28. Brochado, A. R. et al. Species-specific activity of antibacterial drug combinations. Nature. 559, 259–263 (2018).

29. Ragheb, M. N. et al. Inhibiting the Evolution of Antibiotic Resistance. Mol. Cell. 73, 157–165 (2018).

30. Pribis, J. P. et al. Gamblers: An Antibiotic-Induced Evolvable Cell Subpopulation Differentiated by Reactive-Oxygen-Induced General Stress Response. Mol Cell. 74, 785-800.e7 (2019).

31. Rutherford, S.T., Bassler, B.L. Bacterial quorum sensing: its role in virulence and possibilities for its control. Cold Spring Harb Perspect Med. 2: a012427 (2012).

32. Zhu, L. & Lau, G. W. Inhibition of competence development, horizontal gene transfer and virulence in Streptococcus pneumoniae by a modified competence stimulating peptide. PLoS Pathog. 7, e1002241 (2011).

33. Miller, E. L., Evans, B. A., Cornejo, O. E., Roberts, I. S., Rozen, D. E. Pherotype Polymorphism in Streptococcus pneumoniae Has No Obvious Effects on Population Structure and Recombination. Genome Biol Evol. 9, 2546–2559 (2017).

34. Atangcho, L., Navaratna, T., Thurber, G. M. Hitting Undruggable Targets: Viewing Stabilized Peptide Development through the Lens of Quantitative Systems Pharmacology. Trends Biochem Sci. 44, 241–257 (2019).

35. Martin, M. et al. ComE/ComE∼P interplay dictates activation or extinction status of pneumococcal X-state (competence). Mol. Microbiol. 87, 394–411 (2013).

36. Sá-Leão, R. et al. Identification, prevalence and population structure of non-typable Streptococcus pneumoniae in carriage samples isolated from preschoolers attending day-care centres. Microbiology. 152, 367–376 (2006).

37. McGee, L. et al. Nomenclature of major antimicrobial-resistant clones of Streptococcus pneumoniae defined by the pneumococcal molecular epidemiology network. J. Clin. Microbiol. 39, 2565–2571 (2001).

38. Hammerschmidt, S. et al. Illustration of pneumococcal polysaccharide capsule during adherence and invasion of epithelial cells. Infect. Immun. 73, 4653–4667 (2005).

39. Huang, Y. H., Lin, J. S., Ma, J. C., Wang, H. H. Functional Characterization of Triclosan-Resistant Enoyl-acyl-carrier Protein Reductase (FabV) in Pseudomonas aeruginosa. Front Microbiol. 7, 1903 (2016).

40. Heath, R. J., Rock, C. O. Microbiology: A triclosan-resistant bacterial enzyme. Nature. 406, 145–146 (2000).

41. Westfall, C. et al. The Widely Used Antimicrobial Triclosan Induces High Levels of Antibiotic Tolerance In Vitro and Reduces Antibiotic Efficacy up to 100-Fold In Vivo. Antimicrob Agents Chemother. 63: e02312–18 (2019).

42. Martin, B., Prudhomme, M., Alloing, G., Granadel, C., Claverys, J.-P. Cross-regulation of competence pheromone production and export in the early control of transformation in Streptococcus pneumoniae. Mol. Microbiol. 38, 867–878 (2000).

43. Taber, H. W., Mueller, J. P., Miller, P. F., Arrow, A. S. Bacterial uptake of aminoglycoside antibiotics. Microbiol. Rev. 51, 439–457 (1987).

44. Clavé, C & Trombe, M.-C. DNA uptake in competent Streptococcus pneumoniae requires ATP and is regulated by cytoplasmic pH. FEMS Microbiol. Lett. 65, 113–118 (1989).

45. Martín-Galiano, A. J., Ferrándiz, M. J., De La Campa, A. G. The promoter of the operon encoding the F_0_F_1_ ATPase of Streptococcus pneumoniae is inducible by pH. Mol. Microbiol. 41, 1327–1338 (2001).

46. Liu, X. et al. High-throughput CRISPRi phenotyping identifies new essential genes in Streptococcus pneumoniae. Mol. Syst. Biol. 13, 931 (2017).

47. Muñoz, R., García, E., De la Campa, A. G. Quinine specifically inhibits the proteolipid subunit of the F_0_F_1_H^+^-ATPase of Streptococcus pneumoniae. J. Bacteriol. 178, 2455–2458 (1996).

48. Lund, E. & Henrichsen, J. Methods in Microbiology, T. Bergan, J. R. Norris, Eds. (Academic Press, 1978; http://www.sciencedirect.com/science/article/pii/S0580951708703659), vol. 12, pp. 241–262.

49. Lin, D. Y.-W., Huang S, Chen, J. Crystal structures of a polypeptide processing and secretion transporter. Nature. 523, 425–30 (2015).

50. Guiral, S., Hénard, V., Granadel, C., Martin, B., Claverys, J.-P. Inhibition of competence development in Streptococcus pneumoniae by increased basal-level expression of the ComDE two-component regulatory system. Microbiol. Read. Engl. 152, 323–331 (2006).

51. M100Ed28 | Performance Standards for Antimicrobial Susceptibility Testing, 28th Edition. Clin. Lab. Stand. Inst., (available at https://clsi.org/standards/products/microbiology/documents/m100/).

52. Yuzenkova, Y. et al. Control of transcription elongation by GreA determines rate of gene expression in Streptococcus pneumoniae. Nucleic Acids Res. 42, 10987–10999 (2014).

53. Keller, L. E., Rueff, A. S., Kurishima, J., Veening, J.-W. Three New Integration Vectors and Fluorescent Proteins for Use in the Opportunistic Human Pathogen Streptococcus pneumoniae. Genes. 10, 394 (2019).

54. Mosmann, T. Rapid colorimetric assay for cellular growth and survival: application to proliferation and cytotoxicity assays. J. Immunol. Methods. 65, 55–63 (1983).

55. Clementi, E. A., Marks, L. R., Roche-Håkansson, H., Håkansson, A. P. Monitoring Changes in Membrane Polarity, Membrane Integrity, and Intracellular Ion Concentrations in Streptococcus pneumoniae Using Fluorescent Dyes. JoVE J. Vis. Exp. 84, e51008–e51008 (2014).

56. Moreno-Gámez, S. et al. Quorum sensing integrates environmental cues, cell density and cell history to control bacterial competence. Nat. Commun. 8, 854 (2017).

57. Sorg, R. A., Veening, J.-W. Microscale insights into pneumococcal antibiotic mutant selection windows. Nat Commun. 30, 6:8773 (2015).

