## Supplementary material for "Fighting the spread of antibiotic resistance with bacterial competence inhibitors": Materials and Methods, Supplementary Figures 1-7

### *Supplementary methods*

Construction of strain ADP9 (*cep::P₃-luc*).

To verify that the compounds do not influence luciferase acitivty by indirect means, the plasmid pPEP23[17], containing the $P_3$-*luc* construct was transformed into *S. pneumoniae* D39V ($P_3$ promotor drives the constitutive expression of luciferase). Transformants were selected on Columbia blood agar supplemented with 100 µg/ml spectinomycin and correct colonies were verified by PCR and sequencing.

Construction of strain ADP46 (PssbB-ssbB-luc, cps::cat).

The PCR product of the capsular operon deletion, replaced by the chloramphenicol resistance cassette (*cps::cat*), was amplified from ADP25 strain[17] transformed into the MK134 strain[17]. Transformants were selected on Columbia blood agar supplemented with 4.5 µg/ml chloramphenicol and correct colonies were verified by PCR and sequencing.

Construction of strains ADP50 (PMEN18, PssbB-ssbB-luc), and ADP53 (S. sanguinis V683, PssbB-ssbB-luc).

The PCR fragment $P_{ssbB}$-*ssbB-luc* including the kanamycin resistance cassette (MK134 strain), was transformed into the pneumococcal international clone PMEN18, and the reference strain *Streptococcus sanguinis* V683. Transformants were selected on Columbia blood agar supplemented with 250 µg/ml kanamycin and correct colonies were verified by PCR and sequencing.

Construction of strain ADP65 (PssbB-ssbB-luc, prs1::PF6-lacI).

The PCR fragment *prs1::*$P_{F6}$-*lacI* including the gentamycin resistance cassette[45], was transformed into the MK134 strain. Transformants were selected on Columbia blood agar supplemented with 20 µg/ml gentamycin and correct colonies were verified by PCR and sequencing.

Construction of strain ADP73 (pPGs6).

The replicative plasmid pPGs6[52] was transformed to D39V strain. Transformants were selected on Columbia blood agar supplemented with 0.5 µg/ml erythromycin and correct colonies were verified by PCR and sequencing.

Construction of strain ADP148 (bgaA::PssbB-luc, prs1::PF6-lacI, cep::Plac-comCDED58E, comCDE::cat).

To control the production of the phosphorylmimetic C*omE*$^{D58E}$, a strain with an IPTG-inducible *comCDE*$^{D58E}$ was created, with the deletion of the original *comCDE* locus. Using the ADP107 strain as a template (*cep*::$P_{lac}$-*comCDE*), a point mutation in comE gene (T174G) was introduced by overlapping PCR, using primers ADP3/65 (CATGAATATCGATCTCTAGGAAATAAAGC) and ADP3/66 (GCTTTATTTCCTAGAGATCGATATTCATG). This point mutation results in an amino acid replacement (ComE$^{D58E}$). The product was transformed into ADP95, resulting in strain ADP145, and transformants were selected on Columbia blood agar containing 100 µg/ml spectinomycin. Then, the PCR product of Δ*comCDE::cat* from ADP107 strain was transformed into ADP145, resulting in strain ADP148. Transformants were selected on Columbia blood agar containing 4.5 µg/ml chloramphenicol and correct colonies were verified by PCR and sequencing.

Construction of strain ADP272 (bgaA::PssbB-luc-gfp, prs1::PF6-lacI, cep::Plac-comCDT233I, comCDE::cat).

To control the production of ComD$^{T233I}$, which constitutively transphosphorylases ComE, an IPTG-inducible *comCD*$^{T233I}$*E* was created, with the deletion of the original *comCDE* locus. Using the strain ADP107 as template (*cep::*$P_{lac}$-*comCDE*), a point mutation in comD gene (C1719T) was introduced by overlapping PCR, using primers ADP2/19 (GACCAACAATTTCATCTATGTAATTCTGT) and ADP2/20 (ACAGAATTACATAGATGAAATTGTTGGTC). The product was transformed into ADP95 and transformants were selected on Columbia blood agar containing 100 µg/ml spectinomycin, resulting in strain ADP271. Then, the PCR product of Δ*comCDE::cat* from ADP107 strain was transformed into ADP271 and transformants were selected on Columbia blood agar containing 4.5 µg/ml chloramphenicol, resulting in strain ADP272. Correct colonies were verified by PCR and sequencing.

Construction of strains ADP166, ADP168 and ADP172 (PssbB-luc in the CRISPRi strains targeting ATPase-related genes).

To confirm whether pH homeostasis is essential for competence, we used three CRISPRi strains, with sgRNAs targeting three genes of the operon encoding the ATP synthase: *atpC*, *atpA* and *atpE* (strains #342, #349 and #362, respectively[45]. $P_{ssbB}$-*ssbB-luc* fragment including the kanamycin resistance cassette (MK134 strain) was transformed into these three strains, and transformants were selected on Columbia blood agar containing 250 µg/ml kanamycin. Correct colonies were verified by PCR and sequencing.

Construction of strains ADP265, ADP268 and ADP270 (P3-luc in the CRISPRi strains targeting ATPase-related genes).

To determine whether ATPase depletion affects the activity of the luciferase, we transformed the PCR fragment $P_3$-*luc* with integration into the intergenic sequence between gene loci *spd_0422* and *spd_0423* into the above mentioned strains #342, #349 and #362. Transformants were selected on Columbia blood agar containing 100 µg/ml spectinomycin. Correct colonies were verified by PCR and sequencing.

Construction of strains ADP110 (bgaA::PssbB–luc, prs1::PF6-lacI, cep::Plac-comD, ΔcomD) and ADP140 (bgaA::PssbB–luc, prs1::PF6-lacI, cep::Plac-comE, comE::cat).

To examine whether upregulation of *comD or comE* could bypass the COM-block activity of TCL, we constructed a suite of strains in which we could ectopically induce competence genes by the addition of IPTG, with the deletion of the original locus. To amplify *comD*, primers ADP4/37 (CAGTAGATCTAGGAGGAGAGTAATGGATTTATTTGGATTTG) and ADP2/65 (ACGTCTCGAGACTAGTCATTCAAATTCCCTCTTAAATC) were used. To amplify *comE*, primers ADP2/76 (ACTGAGATCTGACAAATCATTAGATTTAAGAGGG) and ADP2/58 (ACGTCTCGAGGCGGCCGCCAATTTCTTGCTAATTGTC) were used, with D39V strain as a template for both fragments. PCR products were digested with BglII and XhoI and were ligated with similarly digested pPEP1 plasmid containing the $P_{lac}$ promoter[45]. The ligations were transformed into strain ADP95 and transformants were selected on Columbia blood agar containing 100 µg/ml spectinomycin, resulting in strains ADP138 and ADP85, respectively. To these strains, either the PCR product *ΔcomD** (clean mutant) or *ΔcomE::cat* was transformed into those strains, resulting in strains ADP110 and ADP140. To construct the ΔcomD* mutant, two STOP codons were introduced in the beginning of the sequence, leaving the polycistronic nature of comDE intact. Overlapping primers were 4/55 (TGGGACGGTTATTGTTCATTAATAAATTATTAGTC) and 4/56 (GACTAATAATTTATTAATGAACAATAACCGTCCC).

To delete *comE*, the original gene was replaced by a chloramphenicol resistance marker. The upstream region was amplified using primers ADP2/65+XhoI (ACGTCTCGAGACTAGTCATTCAAATTCCCTCTTAAATC) and LA11 (GGCGGATCCGGCAGTTTGTGTA), the downstream region with primers pr164+NotI (AGCGCGGCCGCGGGGCTAAATTTAGC) and LA54 (AATCGCCATCTTCCAATCCC), and the chloramphenicol resistance marker with ADP2/79+XhoI (GTAAGGAAATCCATTATGTACCTCGAGCTCGCCCATTAGTTCAACAAACGAAAATTGG) and MT43+NotI (GCATGCGGCCGCGTGACATTAGAAAACCGAC). All three fragments were digested with the proper restriction enzymes (NotI and/or XhoI) and ligated. Transformants were selected on Columbia blood agar containing either no antibiotic (for *Δ*comD* transformation) or 4.5 µg/ml chloramphenicol (from *comE::cat* transformation), and correct colonies were verified by PCR and sequencing.

Construction of strains ADP129 (PcomCDE-comCDE-luc) and ADP137 (PcomAB-comAB-luc).

To test the activity of the early competence promoters driving expression of the *comAB* and *comCDE* operons, we fused the firefly luciferase *luc* to these operons. For ADP129 strain, we PCR-ed the *comCDE-luc* fragment including the kanamycin resistance marker, from strain DJS29[17]. For the *comAB-luc* construct, the upstream region of *comAB* was amplified using primers ADP2/75 (AGGAGAGGATGAAACCAGAATTTTTAG) and ADP2/56+XhoI (ACTGACTAGTCTCGAGCACGAACATTACTCTTTGTTCAA), the downstream region with primers ADP3/33+NotI (CAGTGCGGCCGCTATTTATTCGGTTAAATTCTTGTG) and ADP2/45 (CAGCCTTTTTTTCAACAAAAATACGTTTATC), and the luc gene including the kanamycin resistance marker (from MK134 strain) with ADP4/36+XhoI (CAGTCTCGAGGCTGAAGGAGGAATAATGAGATCCG) and PG97+NotI (CAGTGCGGCCGCTCTAGGTACTAAAACAATTC). All three fragments were digested with the proper restriction enzymes (NotI and/or XhoI) and ligated. For both ligations, transformants were selected on Columbia blood agar containing 250 µg/ml kanamycin, and correct colonies were verified by PCR and sequencing.

Construction of strains ADP279 (bgaA::PssbB-luc, atpEV48L) and ADP280 (bgaA::PssbB-luc, atpEA49T).

To test whether two different point mutations in *atpE* ($AtpE^{V48L}$ and $AtpE^{A49T}$) could cause a loss of susceptibility and/or activity of COM-blockers, we introduced the SNP by overlapping PCR using primers OVL1385 (TGTTTTTAGGTGTTACCTTTATTGAAGGAAC) and OVL1386 (GTTCCTTCAATAAAGGTAACACCTAAAAACA) for $atpE^{A49T}$, and primers OVL1387 (TGTTTTTAGGTCTTGCCTTTATTGAAGGAAC) and OVL1388 (GTTCCTTCAATAAAGGCAAGACCTAAAAACA) for $atpE^{V48L}$, Both fragments were transformed into DLA3, resulting in strains ADP279 and ADP280, respectively. Several transformants were PCR-ed and sequenced to confirm the point mutations.

Construction of strain ADP326 (zip::Plac-HiBiT).

To confirm that HiBiT luminescence is due to the export of the peptide by SPaseI recognition rather than the release by cell lysis, we designed the HiBiT sequence to be controlled by the IPTG-inducible $P_{lac}$ promoter, without the leader peptide. Hence, once HiBiT is produced, is accumulated in the cytoplasm. To do so, we overlap the ADP325 construct with primers OVL2328 (CACCACCACCCATAATAAAATCTCCTTATTTATTTAGATCTTAATTGTGAG) and OVL2329 (CTCACAATTAAGATCTAAATAAATAAGGAGATTTTATTATGGGTGGTGGTG) to remove the leader peptide sequence. Overlapped fragment was transformed into D39V strain and transformants were selected on Columbia blood agar containing 100 µg/ml spectinomycin. Correct colonies were verified by PCR and sequencing.

Construction of strain ADP350 (bgaA::PssbB-luc, prs1::PF6-lacI-tetR, cep::Plac-comAB, comAB::ery, zip::Plac-comCDE, comCDE::cat).

To test wether overexpression of both *comAB* and *comCDE* early operons could overcome the TCL activity, we constructed a strain with both operons under the control of the $P_{lac}$ IPTG-inducible promotor, and the deletion of the native operons. To amplify *comCDE*, primers ADP2/38 (CAGTGGATCCGGTTTTTGTAAGTTAGCTTACAAG) and ADP2/58 (ACGTCTCGAGGCGGCCGCCAATTTCTTGCTAATTGTC) were used, with D39V strain as a template. PCR product was digested with BamHI and XhoI and were ligated with similarly digested pPEPZ plasmid containing the $P_{lac}$ promoter[53]. The ligation was transformed into strain ADP226, and transformants were selected on Columbia blood agar containing 10 µg/ml thrimethroprim, resulting in strain ADP349. Then, *comCDE::cat* fragment from strain ADP107 was transformed into ADP349 resulting in strain ADP350. Transformants were selected on Columbia blood agar containing 4.5 µg/ml chloramphenicol, and correct colonies were verified by PCR and sequencing.

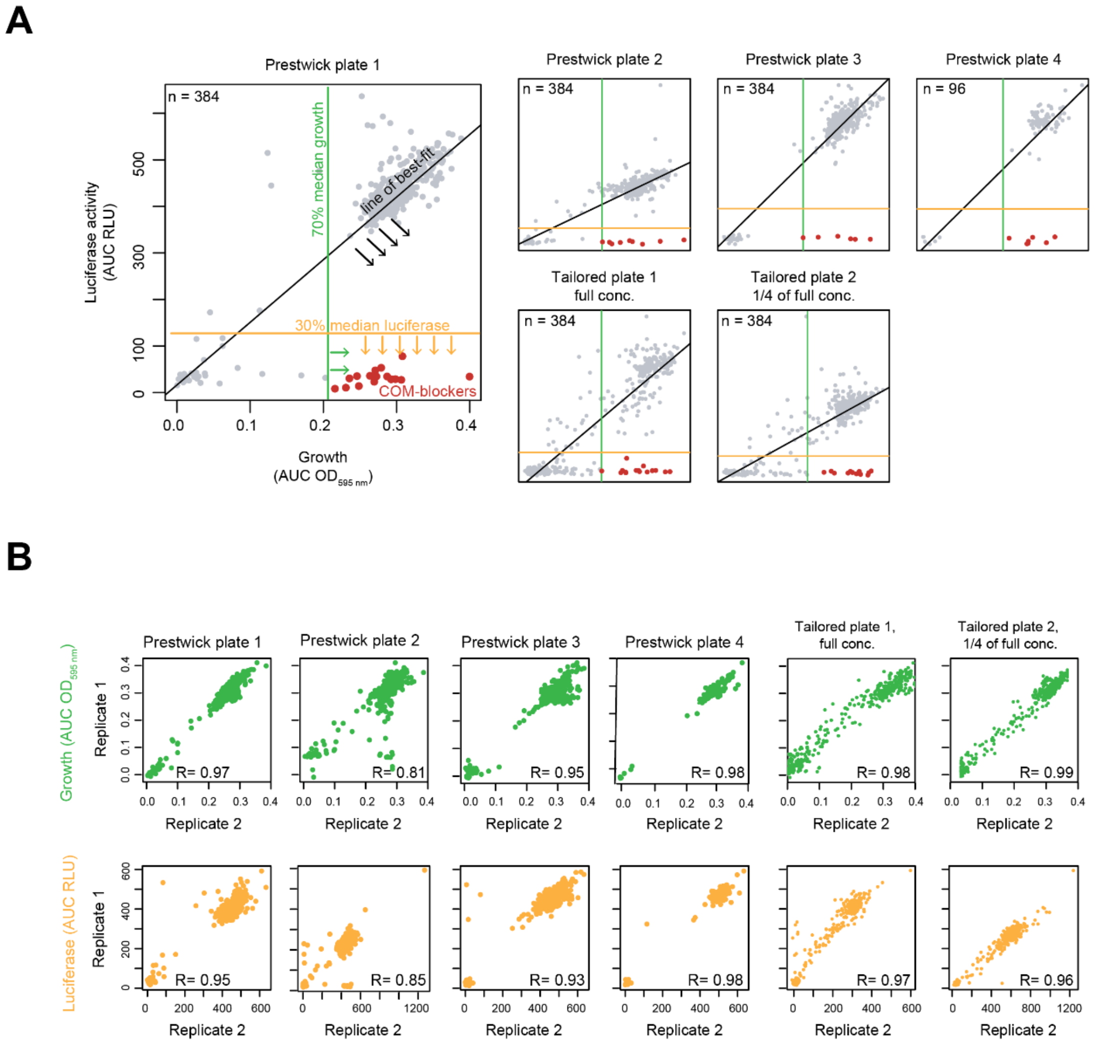


**Fig. S1: Identification of COM-blockers from the high-throughput screen. (A)** Scatter plots show the luminescence signal (area under the curve AUC - RLU) versus growth (AUC - $OD_{595\ nm}$) obtained for each individual well per plate until 5 h of culture (Prestwick library plates 1-4, tailored library 1-2). Only one replicate per plate is shown. The green, yellow and black lines show the stringent criteria used to identify compounds that blocked natural competence development without drastically affecting growth (COM-blockers), here shown in red**. (B)** Replicate correlation for growth and bioluminescence obtained for high-throughput screen. Scatter plots show the correlation between the replicates regarding their growth (green, area under the curve AUC - $OD_{595\ nm}$) as well their luminescence signal (yellow, AUC - RLU), as well their. n indicates the number of drugs (wells) per plate, and R shows that Pearson correlation coefficient.

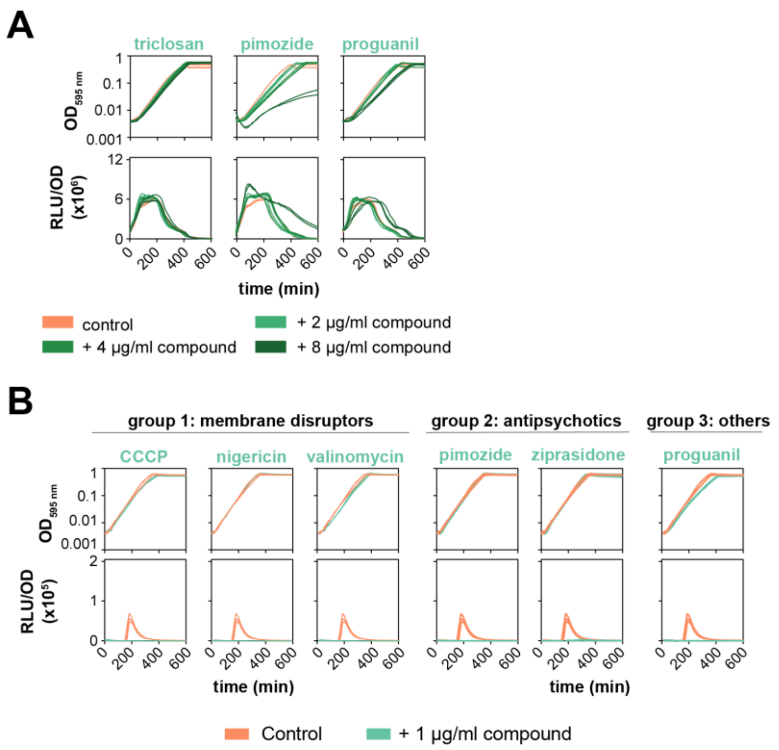


**Fig. S2. Validation of the COM-blockers found in the high-thorughtput screening. (A)** Growth curves and constitutive expression of luciferase in presence of the COM-blockers triclosan, pimozide and proguanil. Strain ADP9 (*cep*::P*3*-*luc*) was grown in C+Y medium at pH 7.5. The addition of the compounds does not affect the luciferase production, discarding a possible interaction between both luciferin and COM-blockers. **(B)** Group 1 compounds that affect the membrane and/or ions homeostasis: cccp, nigericin and valinomycin. The presence of 1 µg/ml of the compound does not affect the growth rates but completely blocks the bioluminescence activity and thereby competence activation. Antipsychotics pimozide and ziprasidone (group 2) and the antimalarial proguanil hydrochloride (group 3) also block competence induction without affecting the growth rates.

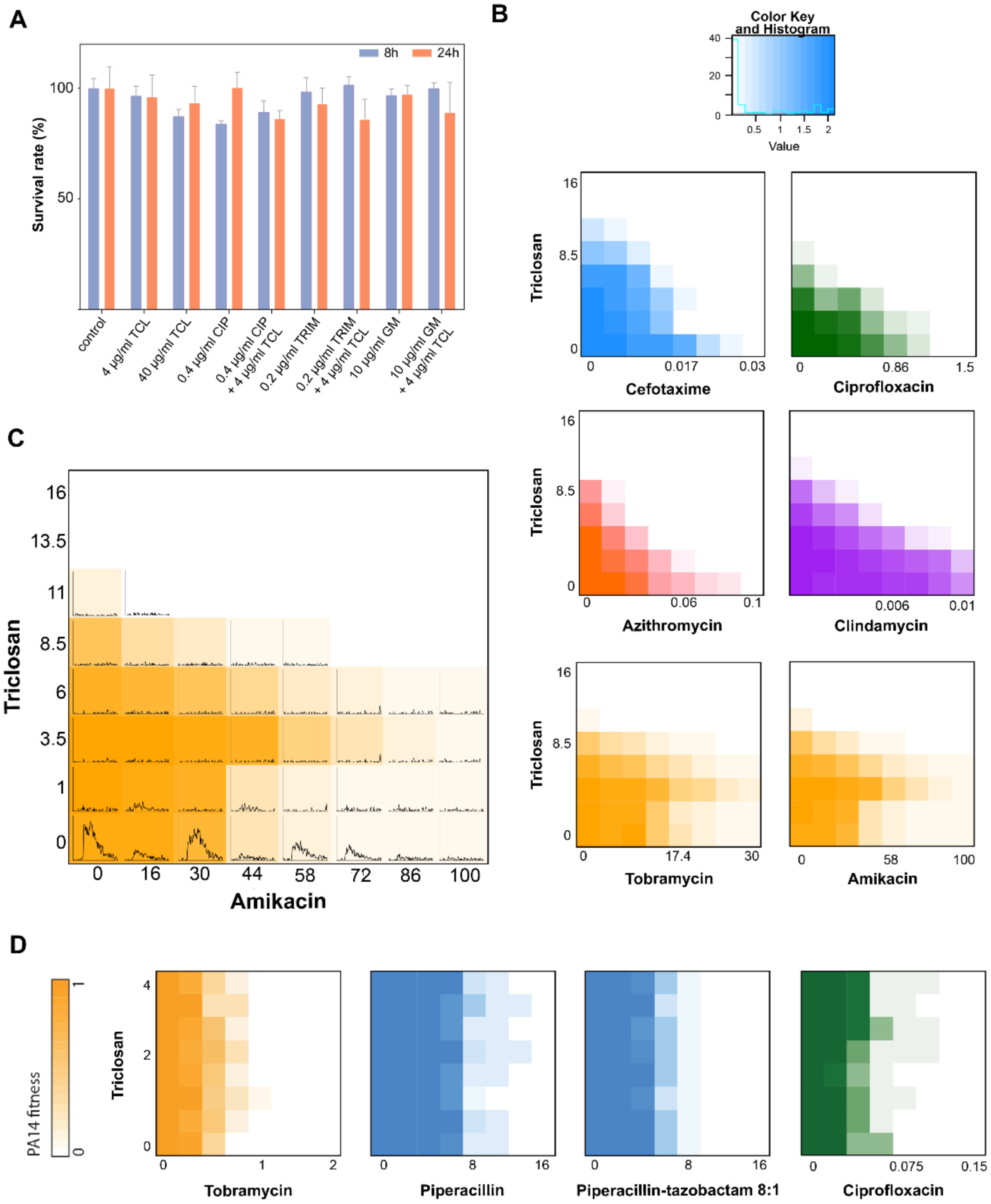


**Fig. S3. Interaction between TCL and antibiotics. (A)** MTT cytotoxicity assay in A549 human cells. The analysis was performed after 8h (blue bars) and 24h (orange bars).The combination of antibiotics plus 4 µg/ml Triclosan does not reduce the viability compared with the antibiotic alone. CIP: Ciprofloxacin, TRIM: trimethoprim and GEN: gentamycin. **(B)** TCL checkerboards with representative antibiotics belonging to the most common classes in *S. pneumoniae*. TCL does not compromise activity of cefotaxime (betalactams class), ciprofloxacin (fluoroquinolones class), azithromycin (macrolides class), clindamycin (lincomycin class). Contrary, a slight interaction was detected with the combination of Triclosan with the aminoglycosides class: tobramycin and amikacin. **(C)** Representation of the competence inhibition by triclosan in the 8x8 checkerboard with the aminoglycoside amikacin (data from panel B). The individual plots in all the growth conditions show the natural competence development (RLU units). Competence was activated only in absence of triclosan. Furthermore, the interaction observed with 3.5 µg/ml of TCL, does not affect the powerful action of this drug. **(D)** Checkerboards of TCL with representative antibiotics in *Pseudomonas aeruginosa*. Addition of TCL does not affect the antibacterial activity of tobramycin (aminoglycosides class), piperacillin (betalactams class) the combination of piperacillin-tazobactam (betalactam + inhibitor of betalactamases) and ciprofloxacin (fluoroquinolones class).

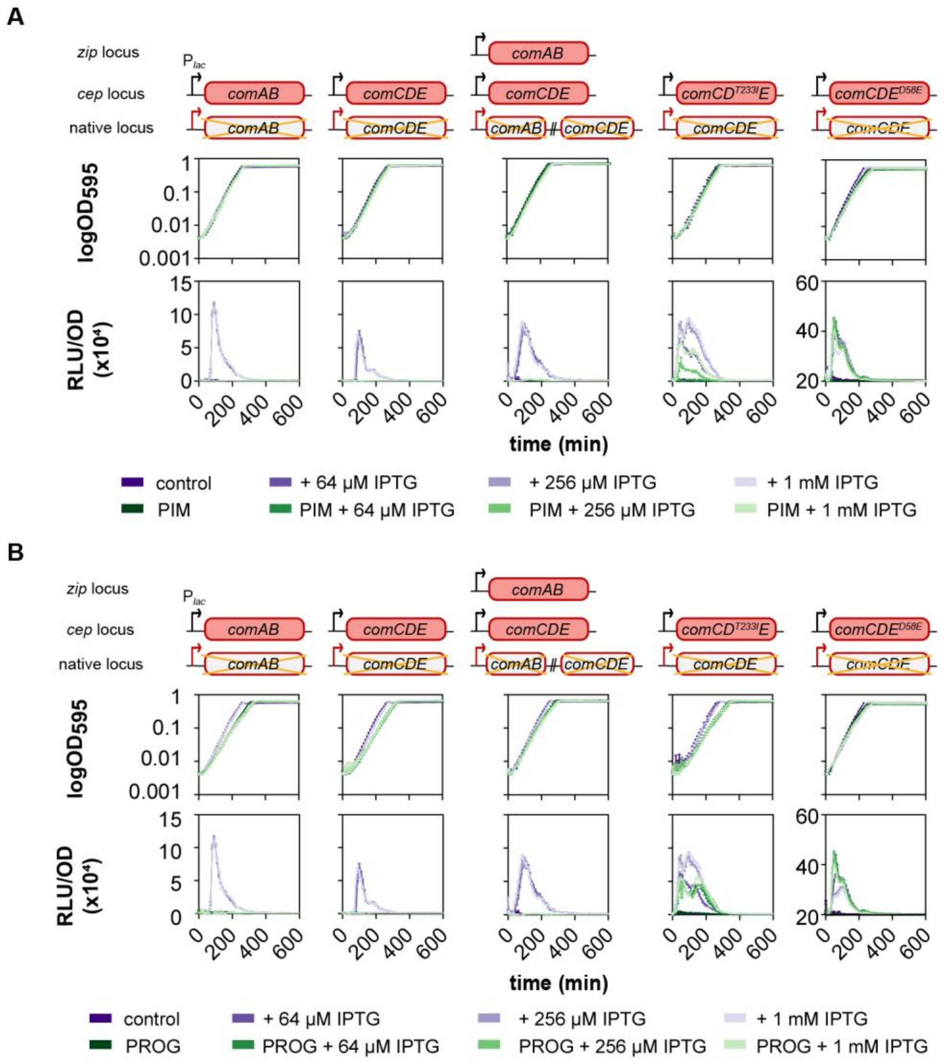


**Fig. S4. COM-blockers counteract the induction of *comAB* and/or *comCDE* and abolished competence. (A)** Pimozide (PIM) effect on growth and detection of competence development. Experiment was performed in C+Y medium at pH 7.5, permissive for natural competence. IPTG was added to the medium at the beginning at different final concentrations. Average of 3 replicates and Standard Error of the Mean (SEM) are plotted. Strains used (left to right: ADP226, ADP107, ADP350, ADP272 and ADP148). Black arrows refer to the IPTG-inducible promoter $P_{lac}$, while red arrows indicate native promoters. (**B**) Proguanil hydrochloride (PROG) effect on growth and detection of competence development. Identical conditions of medium and pH than in panel A. Average of 3 replicates and Standard Error of the Mean (SEM) are plotted. Strains used (left to right: ADP226, ADP107, ADP350, ADP272 and ADP148). Black arrows refer to the IPTG-inducible promoter $P_{lac}$, while red arrows indicate native promoters.

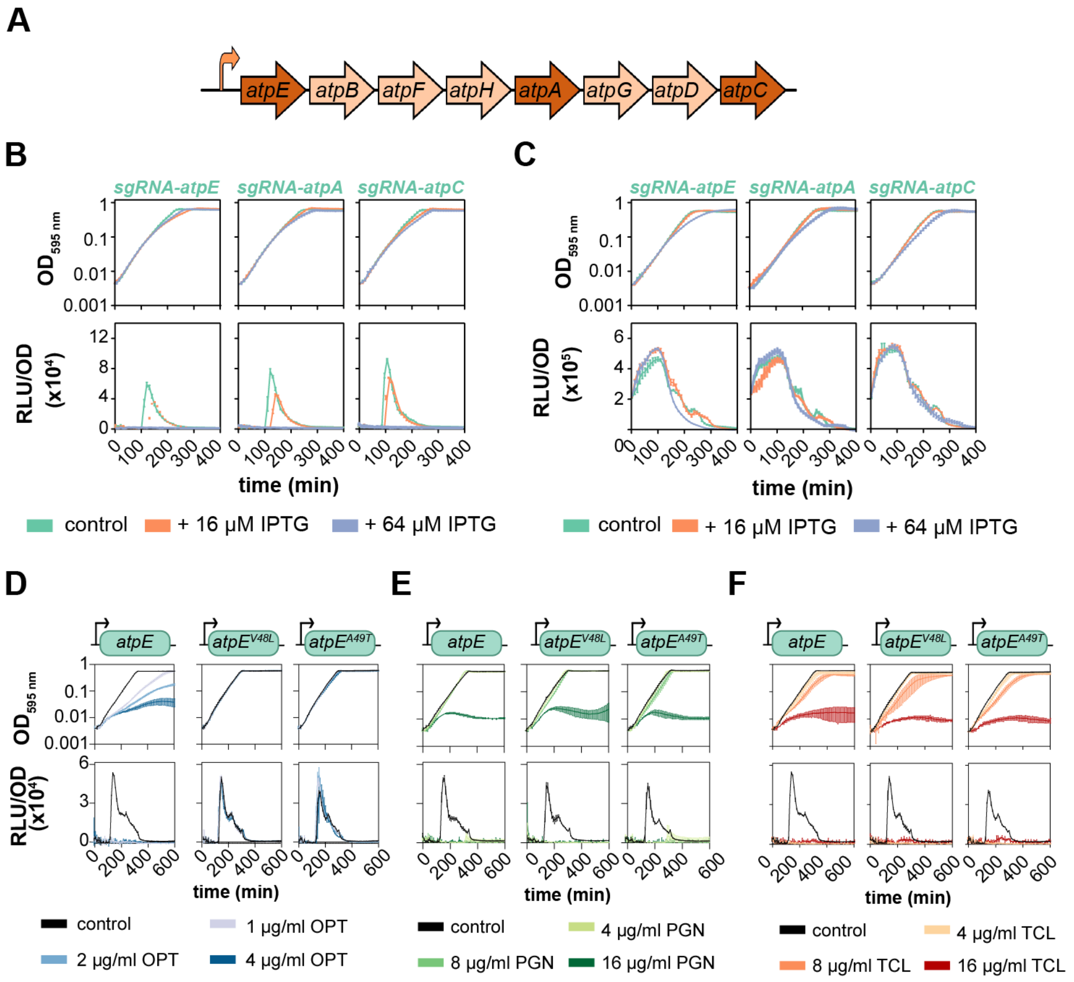


**Fig. S5. $F_0F_1$ ATPase is essential for natural competence induction.** (**A**) Operon encoding for the ATP synthase. Dark color indicates the genes targeted in panels B and C. (**B**) Depletion (by CRISPRi) of gene products related to ATP synthase results in an inhibition of competence (no bioluminescence activity in presence of increasing concentration of IPTG). (**C**) Depletion of these gene products does not affect the luciferase activity in the strains producing constitutive expression of this enzyme. **(D)** Effect of two mutations in AtpE subunit of ATPase of *S. pneumoniae* on optochin activity. Left, growth curves and competence induction of *S. pneumoniae* DLA3 strain without (black) or with a range of optochin. The presence of the compound affects growth rates and competence development. Center and right, the presence of any of both mutations in AtpE (strains ADP279 and ADP280, respectively), restores the growth rates and natural competence development. **(E and F)** Effect of the same mutations on proguanil (PGN, panel E) and TCL (panel F). Both AtpE$^{V48L}$ (strain ADP279) and AtpE$^{A49T}$ (strain ADP280) substitutions have no effect on growth rates or competence inhibition for either proguanil or TCL.

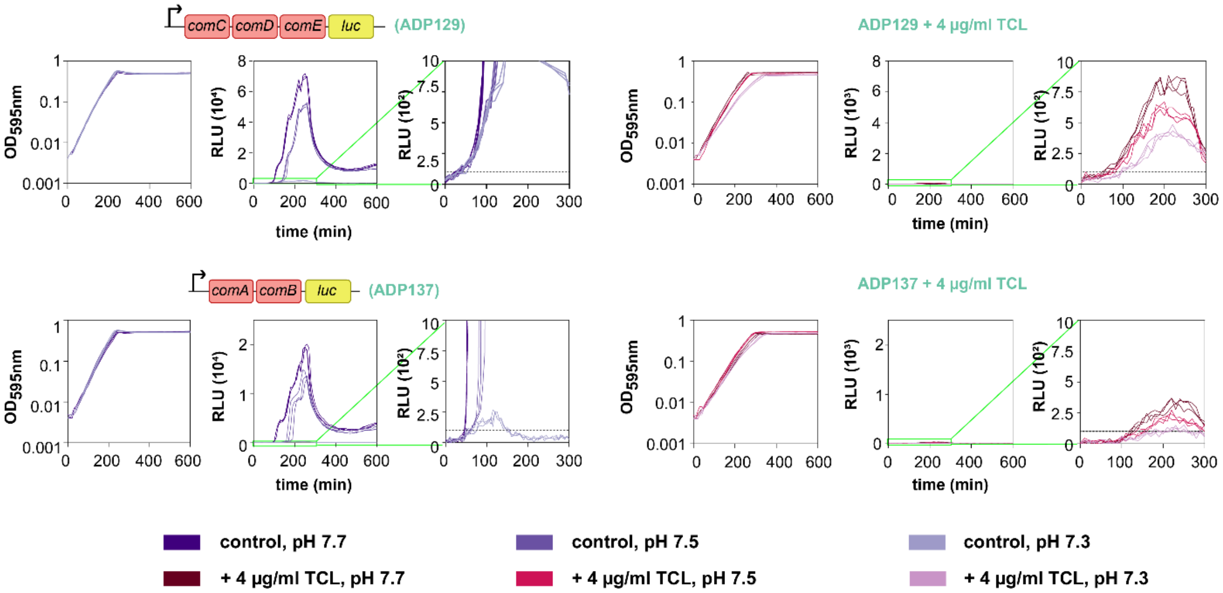


**Fig. S6. Effects of triclosan (TCL) on the activity of the early competence promoters P*comCDE* and P*comAB*.** Strains were grown in C+Y medium in permissive (pH 7.7 and 7.5) and non-permissive (pH 7.3) conditions for natural competence. In normal conditions, there is a basal expression of *comCDE* even at the low pH 7.3. However, the lower production at pH 7.3 results in smaller amounts of CSP accumulated and thereby less phosphorylated ComE (ComE~P) to amplify the competence positive feedback loop[35]. TCL addition nearly completely blocks $P_{comCDE}$ basal expression, which disrupts the positive feedback. Alike, in absence of ComE~P, basal expression of $P_{comAB}$ is shut down in the lower pH and in presence of TCL. Three replicates per each condition are plotted.

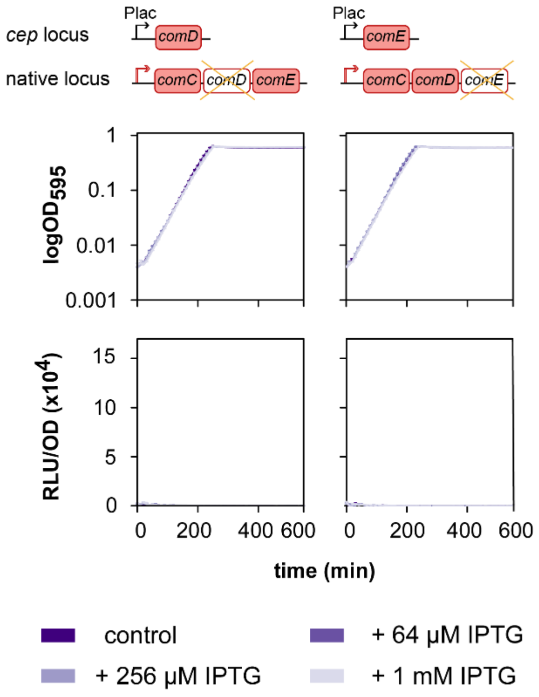


**Fig. S7. Upregulation of comD and comE and effect on competence.** Growth curves and detection of competence development was performed in C+Y medium at pH 7.5, permissive for natural competence. IPTG was added to the medium at the beginning at different final concentrations. Average of 3 replicates and Standard Error of the Mean (SEM) are plotted. Strains used (ADP140 and ADP110, respectively).
